# Oscillatory brain activity as unified control mechanism for working memory and social cognition

**DOI:** 10.1101/2023.02.13.528423

**Authors:** Elisabeth V. C. Friedrich, Yannik Hilla, Elisabeth F. Sterner, Simon S. Ostermeier, Larissa Behnke, Paul Sauseng

## Abstract

It has long been thought that coordination of briefly maintained information (working memory) and higher social cognition (mentalizing) rely on mutually exclusive brain mechanisms. However, here we show that slow rhythmical brain activity in the dorsomedial prefrontal cortex controls distributed networks associated with working memory as well as mentalizing during cognitively demanding visual and social tasks. Depending on the effort necessary for cognitive operations, the phase of slow frontal oscillations is used to precisely tune communication with posterior brain areas. For participants having low autistic personality traits, this mechanism is identical across tasks – no matter whether visual or social information is processed. This underpins a unified function of the mentioned oscillatory brain mechanism in working memory and mentalizing. Participants with high autistic personality traits – thus, with difficulty in social cognition – however, have an inability to efficiently tune brain communication depending on cognitive effort in visual information processing. Even more striking, in higher social cognition they fail to implement coordination of distributed brain networks by slow frontal oscillations completely. While these findings suggest a unified function of brain oscillations in cognitive coordination they also explain why individuals with high autistic personality traits can have difficulties with demanding cognitive processing across domains.

**Significance Statement:** Our findings revealed an interregional brain coupling mechanism based on rhythmical cortical activity to be responsible for successful social and visual working memory by tuning the fronto-parietal network depending on memory load. We suggest that this coupling mechanism can explain how communication between distant brain areas is effectively controlling cognitive functions, independent of the exact type of information that is processed. Importantly, participants with high autistic personality traits struggle with efficient tuning of fronto-parietal networks. Thus, a deficit in this coupling mechanism seems to be an underlying cause of impairments in social and visual working memory, which is often seen in individuals on the Autism Spectrum. These findings might even generalize to other mental disorders as broad cognitive control deficits and social problems are common in a variety of psychiatric and neurological conditions.

---

Imagine you are in charge of assigning the best-suited presents to each of your colleagues during the departmental Christmas party. You will need to infer from your colleagues’ personality traits which type of present they might like; you will have to hold this information in memory and mentally recombine colleague-present pairings. Depending on how well you do on these complex cognitive operations the Christmas party will result in cheers or awkwardness. Processing social information, such as what kind of presents someone might like or dislike, strongly relies on activity within a mentalizing network in the brain ^1–5^. Flexible storage and recombination of information as in the situation described above, however, requires brain activity within a distributed working memory system ^6–14^. Until recently it had been believed that there was an antagonistic relationship between activity within the mentalizing and the working memory networks ^15,16^. Moreover, mentalizing had been investiagted with tasks largely independent of working memory processes ^17,18^. Meyer and colleagues, however, proposed that really effortful social cognition recruits both fronto-parietal mentalizing and working memory networks ^19–21^. Importantly, a key region in coordinating these effortful social cognitive processes seems to be the dorsomedial prefrontal cortex (DMPFC) ^22^.

Brain activity in the dorsomedial prefrontal cortex can reliably be recorded using electroencephalography (EEG). During cognitively demanding processes this is typically manifested by high amplitude, rhythmical EEG activity over fronto-medial recording sites – the so-called frontal-midline theta activity (FM-theta) ^23–26^. Recently, it was demonstrated that this FM-theta activity plays a vital role in coordinating distributed cortical networks during demanding working memory processes ^27,28^. Thereby, fast frequency posterior cortical activity (in the gamma frequency range - which can be considered a proxy for working memory activity ^12,29,30^) is nested into specific phases of the FM-theta cycle ^27^. Berger and colleagues were able to show that in tasks requiring a lot of cognitive control, posterior gamma activity was nested into FM-theta trough – the excitatory phase. Whereas, when a task was relatively easy, posterior gamma was shifted towards 270° and nested more towards the rather inhibitory peak phase. This way, depending on how effortful a working memory process might be and dependent on how much cognitive control is necessary, fronto-parietal cortical networks can either be coupled or dis-engaged. So, if effortful social cognition requires cooperation between a fronto-parietal mentalizing network and a fronto-parietal working memory network, would those networks be coordinated by the above described FM-theta phase mechanism? And if so, would there be interindividual differences in this mechanism as it is known that there is a large variability in terms of mentalizing skills? Individuals with Autism Spectrum Disorder, for instance, do not only display poor mentalizing abilities ^31–33^ but also difficulties with visual (but not so much verbal) working memory ^34–40^. They show difficulties in load modulation, need more cognitive effort, and show deficits especially in highly demanding tasks ^34,41–43^. Moreover, autistic individuals show less fronto-parietal network synchronicity ^44–46^, and autistic traits are correlated with EEG theta activity ^47^.

To test whether FM-theta oscillations are involved in coordination of both mentalizing and working memory networks, and to characterize interindividual differences of this mechanism, we recorded EEG from 100 participants divided into two groups - one group with high (AQ > 20) and one with low autistic traits (AQ ≤ 20) based on the Autism Spectrum Quotient (AQ) ^48^ - while they were doing a social, a visual and a verbal working memory task with two difficulty levels each (Figure 1) ^19^. We expected that in the social as well as in the visual working memory tasks, fronto-parietal networks would be coordinated by FM-theta phase. We hypothesised that the mechanism of posterior brain activity nesting into different FM-theta phase segments depending on cognitive effort (see ^27^) should be observable for the mentalizing network in the same way as for the working memory system in individuals with low autistic personality traits. Participants with high autistic traits were expected to struggle with the social working memory task based on a failure of efficient nesting of posterior high frequency brain activity into the excitatory FM-theta phase. We also hypothesised that participants with high autistic traits would be more rigid in their dynamic control of fronto-parietal networks in the visual working memory task and would show deficits especially in the highly demanding level of the task. These results should therefore provide strong evidence for a general mechanism of controlling working memory as well as mentalizing networks by the means of FM-theta oscillations. And they should explain why cognitive control in higher social cognition as well as working memory are aberrant in individuals with high autistic traits.

**Figure 1.**
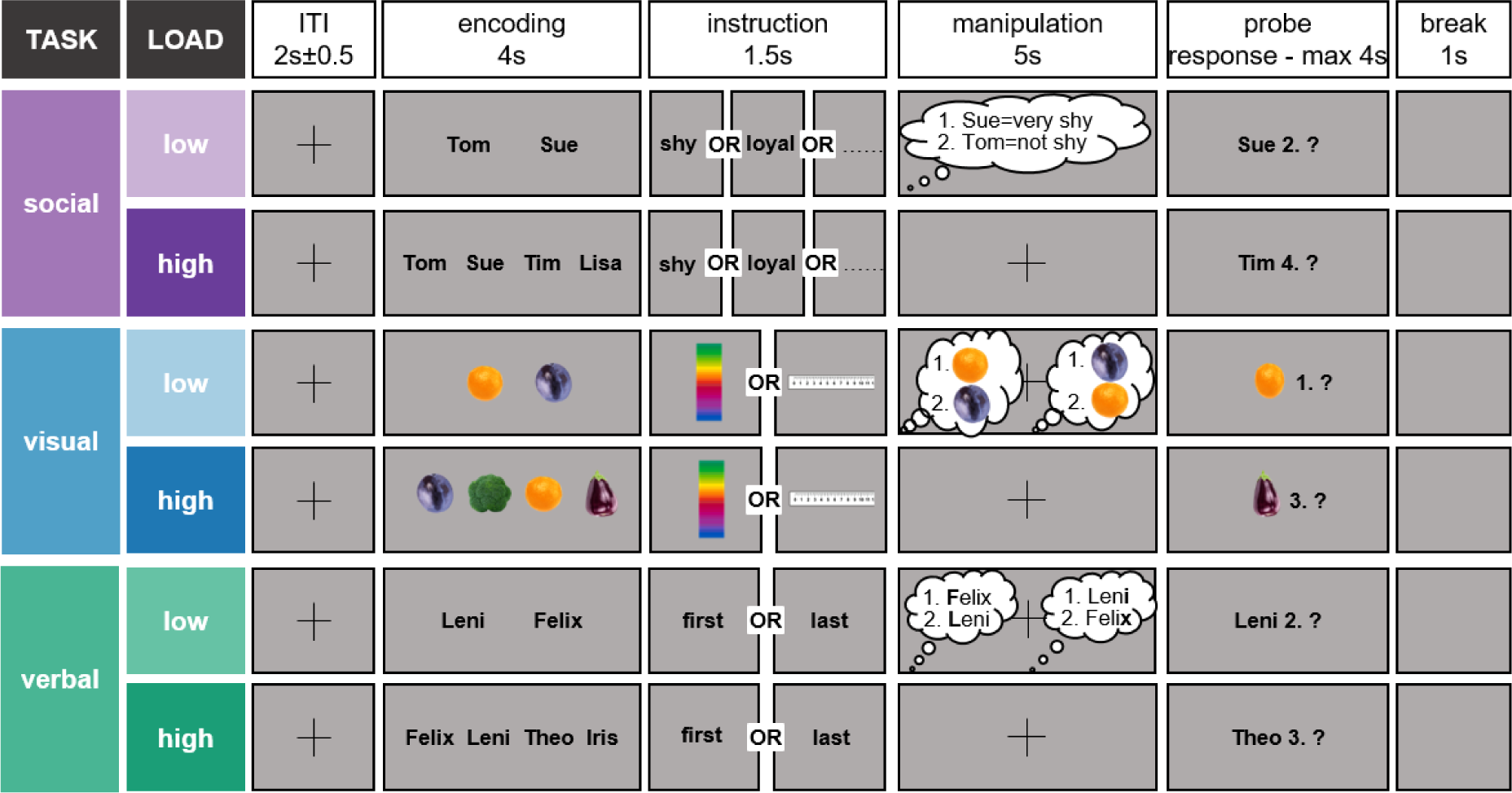
Experimental design. The experiment included three working memory tasks (social, visual and verbal), each consisting of a low and a high load condition. After an inter-trial interval (ITI) jittered around 2 s (± 0.5 s), either two (low load) or four (high load) stimuli were presented for 4 s for the participants to encode. In the social task ^19,21^, the displayed names were drawn from an individualized questionnaire of the screening (i.e., social questionnaire), in which participants rated their personal contacts according to traits. In the visual task, pictures of fruits and vegetables were displayed and in the verbal task, German names were shown. Then the instruction was displayed for 1.5 s: In the social task, participants should mentally rank their personal contacts according to the displayed trait. In the visual task, the objects should be mentally sorted either according to color or size. In the verbal task ^19^, the names should be alphabetically sorted either according to their first or last letter. After the 5-s manipulation period, a probe asked whether a specific stimulus was at a specific position in the mental ranking. The participants pressed yes or no with their right hand as fast and accurately as possible on the left and right mouse key, respectively. Match and non-match probes were randomized and occurred equally often. After the response or maximally after 4 s, there was a 1-s break before the next trial started.

## Results

### Behavioral results

The low autistic-traits group had an AQ of 13.63 (SD = 4.24), whereas the high autistic-traits group had an AQ of 26.61 (SD = 5.21) based on the Autism Spectrum Quotient (AQ) ^48^.

Performance was better (Figure 2A) and faster (Figure 2C) in the low than high load condition across tasks and autistic-traits groups as expected: Participants achieved an average performance of 89.65% (SE = 0.54) in the low load and of 78.27% (SE = 0.69) in the high load condition (Figure 2A; F(1, 92) = 379.19, p < 0.001, partial η^2^ = 0.81). They responded on average after 1149.96 ms (SE = 20.79) in the low load and after 1682.92 ms (SE = 30.13) in the high load condition (Figure 2C; F(1, 92) = 802.17, p < 0.001, partial η^2^ = 0.90). Importantly, this significant load modulation in performance was present in every task and comparable between tasks (Figure 2).

**Figure 2.**
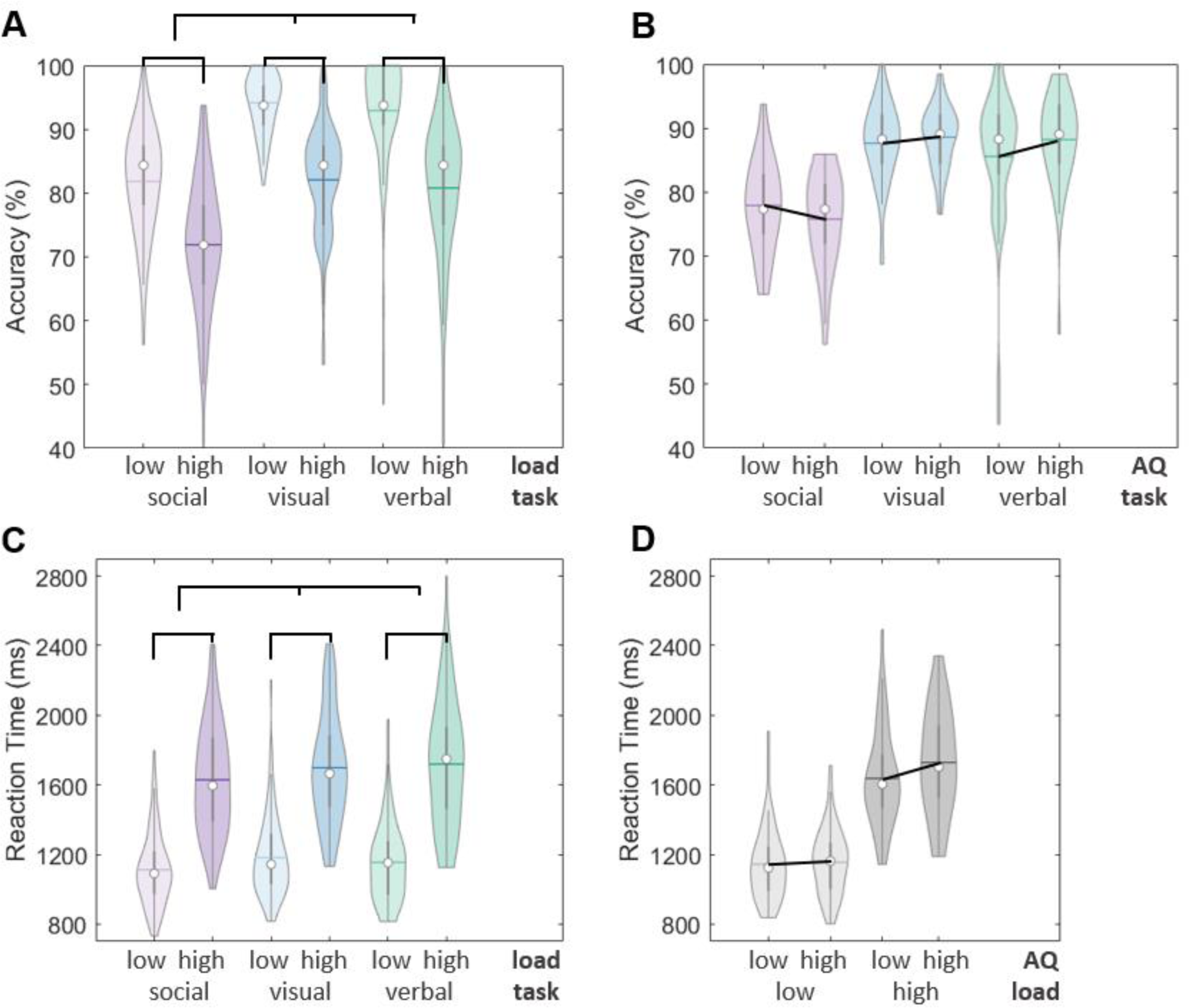
Behavioral results. The data distribution, median (white dot) and mean (colored line) are displayed. **(A)** The low load (light colors) condition showed significantly higher accuracy than the high load (dark colors) condition. The social task (purple) showed significantly lower accuracy than the visual (blue) and verbal (turquoise) tasks. **(B)** There was a significant interaction between tasks and autistic-traits groups in accuracy. **(C)** The participants showed significantly shorter reaction times in the low load condition than the high load condition. They showed significantly shorter reaction times in the social task than in the visual and verbal tasks. **(D)** There was a significant interaction between load conditions and the autistic-traits groups in reaction time.

Accuracy was worse in the social compared to the non-social tasks (Figure 2A; as in ^19^) but reaction times were faster (Figure 2C; as in ^22^): Participants achieved an average performance of 76.86% (SE = 0.74) in the social, 88.13% (SE = 0.57) in the visual and 86.90% (SE = 0.94) in the verbal task (Figure 2A, F(1.67, 153.86) = 90.19, p < 0.001, partial η2 = 0.50). They responded on average after 1371.03 ms (SE = 24.71) in the social, after 1440.62 ms (SE = 25.57) in the visual and after 1437.67 ms (SE = 27.51) in the verbal task (Figure 2C; F(2, 184) = 11.17, p < 0.001, partial η2 = 0.11).

This accuracy-reaction time trade off was reported before in Meyer and colleagues ^19,22^. Despite these differences in performance, Meyer et al.’s fMRI data confirmed the activation of working memory regions in both, social and non-social tasks. The difference in accuracy could be explained by differences in the reliability of the assessment criteria: The criteria for accuracy in the social task (based on an individualized questionnaire of the screening, i.e., the social questionnaire) were not as reliable as in the visual (size and color of vegetables and fruits) and verbal (alphabet) tasks. The re-test reliability of the social questionnaire was on average r = 0.74 (see Table S4). Importantly, the re-test reliability was comparable between autistic-traits groups (Table S4) and thus cannot account for differences between tasks and autistic-traits group in performance reported below.

Besides these main effects, we also found two significant interactions involving the autistic-traits groups:

First, there was a significant interaction between the autistic-traits groups and the load conditions in reaction time (Figure 2D; F(1, 92) = 4.56, p = 0.035, partial η^2^ = 0.05): As expected, individuals with high autistic traits showed similar reaction times in the low load (M_diff_ = 10.95, SE = 41.57, p = 0.793, two-tailed) but a tendency for longer reaction times in the high load condition (M_diff_ = 91.31, SE = 60.27, p = 0.067, one-tailed) in comparison to low autistic-traits participants.

Second, for accuracy a significant interaction between the autistic-traits groups and the tasks was obtained (Figure 2B; F(1.67, 153.86) = 3.46, p = 0.042, partial η^2^ = 0.04): As expected, individuals with high autistic traits showed a tendency to perform worse in the social task (M_diff_ = −2.15, SE = 1.47, p = 0.074, one-tailed) but not in the verbal task (M_diff_ = 2.63, SE = 1.88, p = 0.164, two-tailed) compared to the low autistic-traits group. However, we could not find differences in the visual task in accuracy collapsed over load conditions between the autistic-traits groups (M_diff_ = 0.92, SE = 1.14, p = 0.789, one-tailed). Differences in the visual task between the autistic-traits groups were rather represented by the group differences in reaction time in the high load condition as indicated by the interaction above.

### ROIs and oscillatory FM-theta

We used *a priori* defined coordinates to extract data in source space for the DMPFC as well as for eleven posterior ROIs (see Figure 3B, Table S7; ^19,21,22,49^). Five of these posterior ROIs were reported in literature to be active during social working memory tasks (left/right temporal pole, left/right temporo-parietal junction, medial precuneus) and match with regions from the mentalizing system ^19,21,22^. In previous studies, six of these regions were found to be active in nonsocial tasks or responsible for general load effects in working memory processes (left/right precuneus/posterior cingulate cortex, left/right inferior parietal lobe, left/right intraparietal sulcus) and considered typical working memory regions ^19,22,49^.

**Figure 3.**
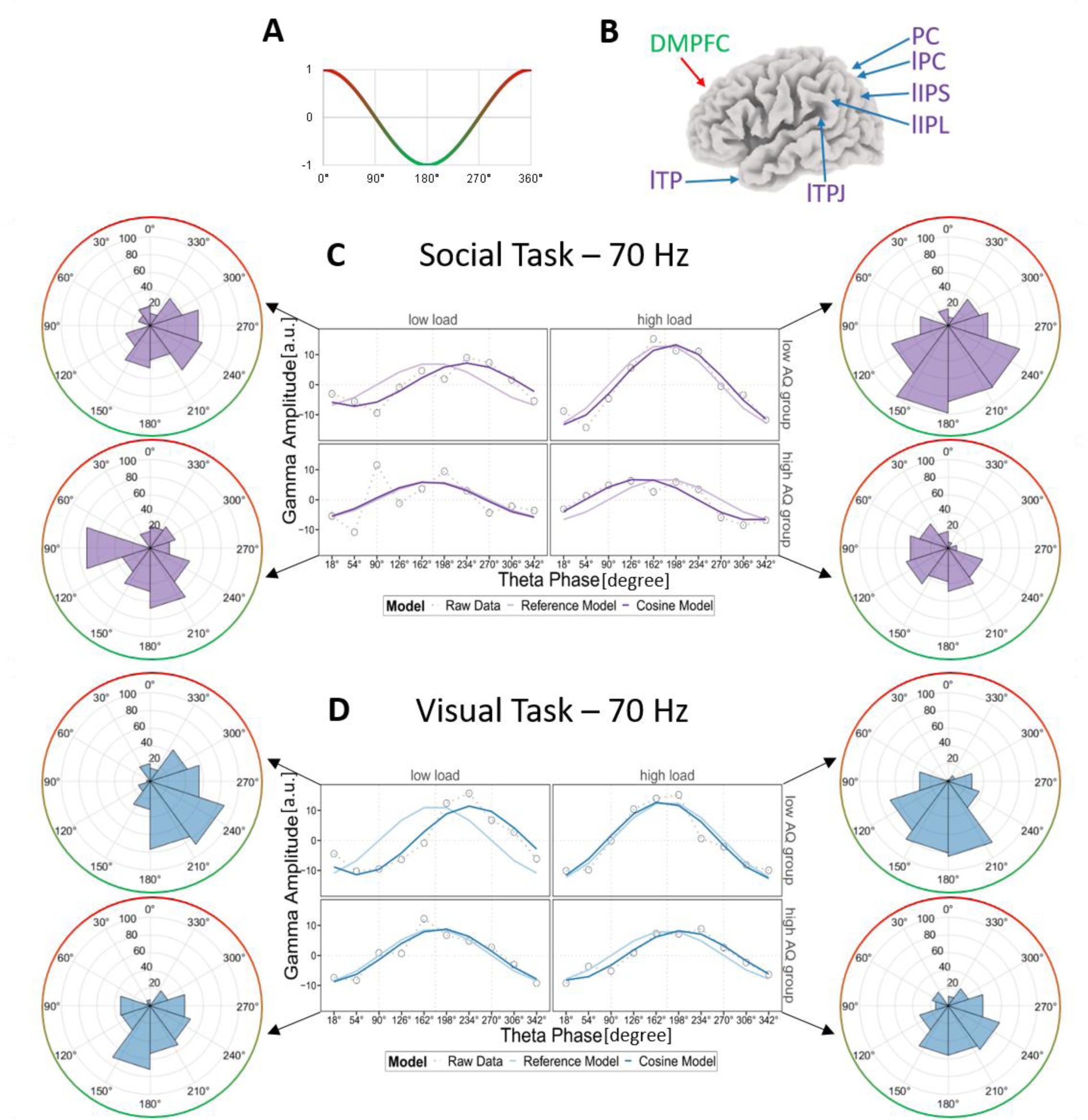
Phase-amplitude coupling. **(A)** The FM-theta trough is at 180° (green) and FM-theta peak is at 0°/360° (red) in our cosine model. **(B)** FM-theta phase was extracted from the dorsomedial prefrontal cortex (DMPFC; green/red). Posterior gamma amplitude (purple/blue) was extracted from 11 posterior regions of interest (ROIs): left/ right temporal pole (l/rTP), left/right temporo-parietal junction (l/rTPJ), left/right inferior parietal lobe (l/rIPL), left/right intraparietal sulcus (l/rIPS), left/right/medial precuneus/posterior cingulate cortex (l/rPC, PC). The arrows show the approximate left and medial ROIs, for all coordinates see Table S7. **(C)** and **(D)** The z-transformed posterior instantaneous gamma amplitude was sorted according to instantaneous FM-theta phase and averaged over all 11 posterior ROIs. In the line charts, the grey dots indicate the empirical z-transformed and sorted 70-Hz gamma amplitudes, the light purple/blue lines our null-shift reference cosine model (simulating that strongest gamma amplitudes were locked in the trough of FM-theta phase) and the dark purple/blue lines the cosine model fitted to our empirical data. In the circular plots, z-transformed sorted gamma amplitudes (blue/purple) are displayed as percentage of the signal, FM-theta peak (0°) is indicated in red and FM-theta trough (180°) in green. In the **(C)** social (purple) and **(D)** visual (blue) tasks in the low autistic-traits (AQ) group (top rows), the highest gamma amplitudes were locked in the FM-theta trough (180°) in the high load (right) and shifted towards 270° in the low load condition (left), demonstrating efficient and load dynamic coordination of fronto-parietal networks. In the **(C)** social task in the high autistic-traits group (bottom row), there was no significant phase-amplitude coupling in the low load and a phase shift towards 90° in the high load condition. This indicates inefficient coordination of working memory and mentalizing networks by FM-theta phase in high autistic-traits individuals. In the **(D)** visual task in the high autistic-traits group (bottom row), the highest gamma amplitudes were locked in the FM-theta trough (180°) already in the low load and slightly shifted in the high load condition, either pointing towards rigidity in working memory control or increased cognitive effort already in the low load condition in high autistic-traits individuals.

The scope of this paper was to investigate theta-phase-to-gamma-amplitude coupling in a hypothesis-driven way. Therefore, first, it was established whether there was oscillatory, periodic FM-theta present in the recorded EEG. We examined the power spectra for the first 2.5 s of the 5-s manipulation period. Participants indicated that they usually had finished their mental ranking within this time window (first half of the manipulation period in the low load condition (see Table S6)). Within the DMPFC, a clear theta peak was visible for the low as well as for the high autistic-traits group (see Figure S3B). For both groups, one peak in the theta range with a power of 0.2 µV^2^ above the aperiodic component (i.e., 1/f like characteristics) was extracted (FOOOF toolbox ^50^, model fit r^2^ > 0.9 (error = 0.2)), confirming a periodic theta oscillation in the DMPFC ^51^. The power spectra of posterior regions did not reveal any theta peak, showing that the theta oscillation was specific to the DMPFC (see Figure S3C).

Extracting the amplitude at the DMPFC for the standard theta frequency band (4-7 Hz; based on ^27^), we could furthermore statistically show that FM-theta amplitude was comparable between the autistic-traits groups (see Table S8). As expected, FM-theta amplitude was higher for the high load than low load condition^27,52^.

### Phase-amplitude coupling

Using complex Morlet wavelet transformation, we extracted FM-theta absolute phase values for 4-7 Hz from the in source space *a priori* defined DMPFC as well as gamma amplitude between 30-70 Hz in 5 frequency bands from *a priori* defined posterior ROIs (see Figure 3B, Table S7; ^19,21,22,49^) for the first 2.5 s of the 5-s manipulation period. Posterior gamma amplitudes were then z-transformed for each trial, all trials were concatenated, and instantaneous gamma amplitude values from all posterior ROIs were then sorted according to the instantaneous FM-theta phase of the DMPFC. This resulted in sorted posterior gamma amplitude as a function of FM-theta phase. If gamma amplitude is equally distributed across FM-theta phases this will indicate no association between gamma amplitude and FM-theta phase. If, however, sorted gamma amplitude varies across different FM-theta phases, it will suggest interaction between FM-theta and posterior gamma (see ^27^ for details and Figure S4). The sorted posterior gamma amplitude values were used as dependent variable in regression models for the different working memory tasks with FM-theta phase segments, load, posterior ROIs and autistic-traits groups as predictor variables.

Regression models on FM-theta sorted gamma amplitude indicated significant interactions between load and autistic-traits group for the social task (at 70 Hz: AICc = 34.51; at 60 Hz: AICc = 20.84) and the visual task (70 Hz: AICc = 17.66). These effects generalized across posterior ROIs. There was no effect for posterior ROIs (AICc = −109.90 to 2.99, n.s.), posterior ROIs x load (AICc = −249.16 to −131.95, n.s.), posterior ROIs x autistic-traits group (AICc = −232.44 to −126.13, n.s.) or posterior ROIs x load x autistic-traits group (AICc = - 500.64 to −318.89, n.s.) in any of the gamma frequency bands. This result indicates that gamma amplitude at all posterior ROIs (i.e., from the mentalizing network as well as those from the working memory system) was similarly modulated by FM-theta phase. This is why for any further analyses, sorted gamma amplitudes across all posterior ROIs were averaged. Regression analysis for the verbal task did not yield any significant effects (AICc = −28.38 to 8.81, n.s.).

It is important to note that the reported effects of FM-theta phase-sorted gamma amplitude (see above) were not driven by any general gamma amplitude differences between autistic-traits groups or load condition (see Table S8). We also did not obtain any significant interactions involving posterior ROIs with load or autistic-traits group for the 60- and 70-Hz gamma band in the social and visual tasks (see Table S8). Moreover, gamma amplitudes were z-transformed for the phase-amplitude coupling, thus equalizing any possible absolute amplitude differences.

In order to further characterize the nature of the significant modulation of FM-theta phase-sorted gamma amplitude, we fitted our sorted gamma amplitude values to a theta cosine model. A good model fit indicates that posterior gamma amplitudes are periodically modulated by FM-theta phase. We computed the mean absolute error (MAE; the lower the value the better the fit) in order to determine how well our empirical data fit to a cosine model. This cosine model was then compared (1) to an intercept model to check if the modulation of gamma amplitude was significant, separately for respective load condition and autistic-traits group (AIC model comparison cosine vs. intercept); and the cosine model was compared to (2) a null-shift reference cosine model with the trough at 180° to establish whether posterior gamma was precisely locked to the FM-theta trough or to any other FM-theta phase segment.

### Low autistic-traits participants show expected phase-amplitude coupling

Low autistic-traits individuals’ posterior 70-Hz gamma amplitude was periodically modulated by FM-theta phase, both under low and high load conditions in the social and visual tasks (Figure 3C and D top rows). In support of this, their averaged posterior 70-Hz gamma amplitude values barely deviated from the predicted cosine model values as indicated by a very small MAE (social task: MAE_low load_ = 1.67; MAE_high load_ = 2.40; visual task: MAE_low load_ = 2.17; MAE_high load_ = 2.39). Moreover, their data significantly fit a cosine model better than an intercept model (social task: AICc_low load_ = 6.11; AICC_high load_ = 15.16; visual task: AICc_low load_ = 10.68; AICc_high load_ = 14.68). In the low load condition, strongest gamma activity was significantly shifted towards 270° by on average 53.57° (CI = [27.57°, 79.57°]) in the social and 57.39° (CI = [37.54°, 77.20°]) in the visual task. In the high load condition, there was neither a significant phase shift in the social (CI = [−2.37°; 28.57°]) nor visual task (CI = [−23.18°; 8.58°]). This indicates that the maximal gamma amplitude was precisely locked to the FM-theta trough (180°) in the low load condition. In the 60-Hz frequency band in the social task, the phase-amplitude coupling was similar but slightly weaker than in the described 70-Hz frequency band (see Figure S5).

### High autistic-traits participants show a deviant brain communication pattern

In the visual task (Figure 3D bottom row), high autistic-traits individuals’ posterior 70 Hz gamma amplitude was periodically modulated by FM-theta phase both in low (MAE = 1.61, AICc = 11.28) and high load conditions (MAE = 4.12, AICc = 17.25). However, there was no significant phase shift in the low load condition (CI = [−9.89°, 28.45°]).

In the high load condition of the visual task, high autistic-trait individuals’ posterior gamma was slightly shifted away from the FM-theta trough (minimal but significant phase shift of 23.30°; CI = [9.47°, 37.11°]).

In the social task (Figure 3C bottom row), in the low load condition, the high autistic-traits individuals’ data better fit to the intercept model (AICc = −4.88, MAE = 5.17), suggesting no significant periodically modulation of 70-Hz posterior gamma amplitude by FM-theta at all. In the high load condition, posterior gamma amplitude was periodically modulated (AICc = 8.88, MAE = 5.84). However, the maximum of sorted gamma amplitude was shifted towards 90° by 37.39° (CI = [−59.40°, −15.38°]) on average. A shift towards 90° (in contrast to a shift towards 270° as expected based on ^27^), might point to suboptimal communication of information in the fronto-parietal network. Again, phase-amplitude coupling was similar in the 60-Hz frequency band (see Figure S5).

### Circular-linear correlations between phase-amplitude coupling and personality traits

We ran circular-linear correlations between individual instantaneous FM-theta phase and autistic-trait measures (AQ subscales) across participants. Instantaneous FM-theta phase co-occurring with the maximum of sorted 60-Hz frequency band amplitude in the social task was significantly correlated with the AQ subscales imagination in the low load condition (r = 0.30, p = 0.014) and with social skills in the high load condition (r = 0.26, p = 0.036).

## Discussion

This study’s findings indicate that posterior working memory and mentalizing regions are co-activated and controlled by the identical prefrontal, oscillatory mechanism during social as well as visual working memory tasks: posterior gamma amplitude is modulated by FM-theta phase so that optimal time windows for fronto-parietal neural communications arise (see ^27^ for comparison). This brain coupling pattern is modulated by task load; more effortful processing leading to more precise modulation of posterior gamma activity by FM-theta phase. Moreover, our results indicate that individuals with high autistic personality traits appear to have difficulties dynamically adjusting fronto-parietal connectivity dependent on task demands – particularly in social tasks.

Importantly, these findings were based on solid ground as our study included 100 participants and was conducted hypothesis-driven with *a prior* defined ROIs and frequency bands. We could replicate load modulation in performance comparable between tasks ^19,22^ and previous phase-amplitude coupling findings in the visual domain ^27^. Our control analysis for FM-theta and posterior gamma amplitude confirm that differences between the autistic-traits groups really do rely on the phase-amplitude coupling mechanism and cannot be explained by mere differences in amplitude, signal-to-noise ratio and accuracy of phase estimates ^51^.

### Working memory and mentalizing networks unified in low autistic-traits participants

In the low autistic-traits group, posterior high frequency bursts were locked close to the trough of the FM-theta cycle in the high load and shifted towards 270° of the FM-theta cycle in the low load condition in both the social *and* the visual tasks. This is exactly replicating a finding previously reported in young, healthy participants during a visuo-spatial working memory task ^27^. The current results clearly show that FM-theta phase plays a general and vital role in coordinating distributed cortical networks during higher cognition such as working memory processes, largely independent of the specific modality (i.e. visual as well as social).Not only was the factor posterior ROI not significantly involved in any interactions in the regression analysis – indicating that posterior working memory and mentalizing regions displayed a very similar phase-amplitude coupling pattern – an exploratory, supplemental analysis with the working memory and the mentalizing network displayed separately (Figure S6) also clearly suggests that those posterior clusters of regions responded nearly identical to each other in low autistic-traits individuals. Thus, FM-theta phase does not only seem to control the fronto-parietal working memory system but also the mentalizing network. As expected, the DMPFC constitutes the core region linking both systems ^2,19,22,27,53–55^, leading to our proposition of shared neural bases between the visual and social tasks.

In contrast, the verbal task did not elicit the same phase-amplitude coupling in the group with low autistic traits as seen in the other two tasks. The verbal working memory task might rely more strongly on the left dorsolateral prefrontal cortex (lDLPFC) ^19^, which was associated with general working memory functions across verbal, visual, social and emotional working memory tasks in earlier neuroimaging studies ^6–8,14,19,54,56–58^. Thus, we analyzed our data with exactly the same procedure but used the left DLPFC instead of the DMPFC for theta phase extraction (see Table S9). Regression models with theta phase segments from the left DLPFC, load and autistic-traits group as interaction terms reached significance for 50-Hz posterior gamma amplitude in the verbal (AICc = 40.40) and visual (AICc = 28.07) tasks but not in the social task (AICc = −14.79 to −1.75, n.s.). However, the anticipated phase-amplitude coupling (as obtained with DMPFC theta activity for individuals with low autistic traits), could not be systematically found for the left DLPFC (see Figure S7 and S8).

So, in individuals with low autistic traits we found a unified mechanism coordinating both mentalizing and visual working memory processes: FM-theta from the DMPFC coordinated posterior regions independently of whether they were rather considered as part of the working memory or mentalizing network, incorporating effortful social cognition and demanding visual working memory tasks.

### Phase-amplitude coupling explains differences in autistic-traits groups

As discussed above, only in the high load condition, low autistic-trait individuals tuned their fronto-posterior networks towards maximally efficient communication by precisely nesting posterior gamma into the trough of FM-theta (see ^27^ for a discussion). The low load condition demanded less deployment of cognitive resources and less cognitive control, which meant that low autistic-trait individuals did not need to precisely nest posterior gamma into the FM-theta trough but could allow for a slight theta phase shift towards 270°. This dynamic regulation of cognitive control did not optimally function in high autistic-trait individuals. Already in the low load condition of the visual task, they displayed ultimate precision of FM-theta phase to gamma coupling; meaning that they cannot further increase cognitive control when the task becomes more difficult (high load condition).

This pattern might either reflect a certain rigidity of cognitive control functions, i.e. a deficit in load modulation ^43,59,60^, resulting in high autistic-traits individuals being stuck in the effortful mode and being unable to efficiently couple and decouple the fronto-parietal networks dependent on load (Figure 3).

Alternatively this finding could indicate the expenditure of more cognitive effort to achieve comparable performance as the low autistic-traits group ^36,61–63^. An observation rather speaking for the latter interpretation is that in the high load condition of the visual task, high autistic-trait individuals’ posterior gamma was slightly shifted away from the FM-theta trough (Figure 3). This might indicate that their optimal level of cognitive demand is closer to the low load condition, and that there is no optimal compensation mechanism for the high load condition ^34,41,42^. One might speculate that this could support the idea of an inverted U shape of working memory regions activation depending on load ^64^. Our assumptions are also in line with our behavioral data, showing similar reaction times in the low load but a tendency for longer reaction times in the high load condition for high autistic-traits in comparison to low autistic-traits participants (Figure 2D).

In the social task, individuals with high autistic traits failed to show efficient coordination of fronto-parietal networks by FM-theta phase to gamma coupling (Figure 3).

These findings of aberrant phase-amplitude coupling are in line with studies indicating that individuals with Autism Spectrum Disorder display lower activity in the prefrontal cortex and dysfunctional long-rang connectivity, particularly frontal-posterior disconnectivity at lower frequencies ^44–46^. Our behavioral data suggesting the high autistic-traits group tending to perform worse than the low autistic-traits group in the social task, which requires reasoning about others ^31–33^ (see Figure 2B) are also well in line with the described pattern of reduced FM-theta phase to gamma amplitude coupling in the social task.

So, while phase-amplitude coupling is aberrant in the group with high autistic traits, this is also reflected in the behavioral data. Might FM-theta phase to gamma amplitude coupling, therefore, be a general marker of autistic personality traits?

Significant circular-linear correlations between individual instantaneous FM-theta phase and autistic-trait measures (AQ subscales) suggest so. We found that our phase coupling mechanism in the social task was significantly correlated with the AQ subscales imagination in the low load condition and with social skills in the high load condition. The AQ subscale imagination evaluates if people find it easy to create a picture in their mind or play pretend games. Mental imagery can also be associated with visual working memory processes ^65,66^. The AQ subscale social skills evaluates if a person seeks social encounters and manages them well. These correlations highlight the relevance of theta oscillations for autistic traits ^47,67^ and demonstrate that phase-amplitude coupling is functionally relevant. Moreover, they can explain why individuals with high autistic traits might have problems in higher social cognition as well as visual working memory.

Importantly, the reported differences in phase-amplitude coupling and behavior were based on mostly subclinical individuals. As Autism is a spectrum disorder, it is hard to set a clean cut-off between neuro-typical individuals and individuals with a disorder. Although this study included a wide range of participants in respect to the Autism spectrum, they were comparable in age, gender, education and intelligence. Strikingly, differences in the extend of autistic traits-while controlling for other confounding varibales-were enough to manifest themselves in different patterns of efficient coupling and decoupling of fronto-parietal networks. Future studies shoud take the developmental component into account ^43^. This phase-amplitude mechanism might serve as an early predictor for aberrant development. Moreover, it might also help to understand deficits in other mental disorders such as schizophrenia better ^68^.

## Conclusion

In this study, we were able to show that FM-theta at DMPFC coordinates the mentalizing network as well as the working memory system by dynamically nesting posterior brain activity into different FM-theta phases in effortful social and visual working memory tasks. While the visual and social working memory tasks share this neural mechanism, the verbal domain seems to involve different control mechanisms.

Moreover, we demonstrated that interindividual differences in autistic traits influenced efficient coordination of fronto-parietal regions in both networks, the mentalizing and working memory system. Whereas load-dependent and dynamic allocation of resources were seen in the low autistic-traits group, individuals with high autistic personality traits remained in a high effort mode of fronto-parietal network control in the visual task and failed to efficiently coordinate distributed networks by FM-theta phase entirely in the social task. This phase-amplitude coupling mechanism was functionally relevant and could explain deficits in mentalizing and visual working memory in individuals with high autistic traits.

## Methods

### Sample size

We based our sample size on an *a priori* power analysis (MorePower 6.0.4. ^69^) for a task x load repeated-measures ANOVA with autistic-traits group as between-subject factor using an alpha level of 0.05, statistical power of 0.9 and an effect size of partial η^2^ = 0.07. This effect size was a conservative estimation based on both, relevant behavioral(effect size of η^2^ ≥ 0.08 for behavioral performance in the social and verbal tasks ^19^) as well as EEG findings (partial η^2^ = 0.1 for phase-amplitude coupling in the visual task ^27^). We added roughly 15% to the suggested 88 participants to compensate for possible exclusions and thus, recorded data of a total of 100 participants.

### Screening

Participants were either assigned to the low or to the high autistic-traits group according to their Autism Spectrum Quotient (AQ) ^48^. It assesses social skills, attention switching, attention to detail, communication and imagination in five subscales. The scale is sensitive enough to measure autistic traits in the non-clinical population and correlates with social and visual cognitive performance and neurophysiological measures ^35,48,70,71^.

We chose a cut-off of 20 to discriminate between sub-clinical individuals with low autistic traits (i.e., AQ ≤ 20: typical control participants) and high autistic traits (i.e., AQ > 20: individuals who show some difficulties in social cognition such as was shown for relatives of individuals with Autism Spectrum Disorder or specialists in mathematics for example) ^48,72,73^.

In addition to the AQ, all participants underwent a two-step screening process (see Table S1). In the first step, we asked about sociodemographic data and performed a neuropsychiatric telephone interview to only include participants who had no clinically significant neurological or psychiatric disorder. We included individuals with scores above 32 on the AQ ^48^ or a diagnosis of Autism Spectrum Disorder as long as participants were not significantly impaired in their daily-life functioning. The second step was an in-person screening which took place on average 14 days prior to the EEG recording. Handedness was confirmed with the short form of the Edinburgh Handedness Inventory ^74^ and color vision with the Ishihara color test. Verbal and non-verbal intelligence was assessed with the Multiple Choice Vocabulary Test (MWT-B) ^75^ and the Culture Fair Intelligence Test (CFT 20-R, Scale 2) ^76^, respectively. The German short version of the Big Five Inventory (BFI-K) was applied to evaluate the Big Five personality traits ^77^.

Finally, participants filled out the social questionnaire ^19,21,78^. They named 10 personal contacts and evaluated them on 48 German traits on a 1-100 rating scale in intervals of 5 points (see Table S2 and S3) ^79,80^. Personal contacts which were ranked ≥ 15 points apart from one another for a specific trait were used to create individualized trials for the social task in the EEG experiment.

### Participants

Of the 100 volunteers, 51 belonged to the low autistic-traits group (AQ ≤ 20; M_AQ_ = 13.63, SD_AQ_ = 4.24; 25 female and 26 male) and 49 to the high autistic-traits group (AQ > 20; M_AQ_ = 26.61, SD_AQ_ = 5.21; 27 female and 22 male). Both groups were on average 24 years old (range between 18-39 years; SD = 4.42 for the low and 3.84 for the high autistic-traits group). All volunteers had at least a secondary school diploma, German as their native language, were right-handed and had normal or corrected-to-normal vision. None of them was pregnant or had any clinically significant neurological or psychiatric disorder that led to significant impairment in daily-life functioning. Three participants met the criteria for Autism Spectrum Disorder. Please see Table S1 for more details about the participants. There was no difference between the autistic-traits groups in respect to their verbal and non-verbal intelligence scores (CFT 20-R: N_lowAQ_ = 51, M_low AQ_ = 114.27, SE_low AQ_ = 1.85 vs. N_highAQ_ = 49, M_high AQ_ = 118.45, SE_high AQ_ = 1.88, t(98) = −1.58, p = 0.116, two-tailed; MWT: N_lowAQ_ = 51, M_low AQ_ = 103.71, SE_low AQ_ = 1.61 vs. N_highAQ_ = 49, M_high AQ_ = 106.08, SE_high AQ_ = 1.40, t(98) = −1.11, p = 0.270, two-tailed). Participants were mainly recruited within the LMU Munich and Technical University of Munich student communities and were compensated with course credit or 10 € per hour. All participants gave written informed consent prior to participation in the study, which was approved by the LMU Munich Faculty 11 ethics committee.

### Experimental design

The experimental design included a social, visual and verbal working memory task, each implemented in a low (i.e., two stimuli) and a high load condition (i.e., four stimuli to be encoded and manipulated) (Figure 1 and for more details Figure S1, Table S2, S3 and S5). The social and verbal tasks were adapted from Meyer and colleagues ^19,21^.

In the social task, the names of the personal contacts - which were indicated in the social questionnaire during the screening process - were used as stimuli. After encoding the names for 4 s, a trait was presented for 1.5 s (e.g., shy), which was also drawn from the social questionnaire. During the 5-s manipulation period, participants mentally ranked their contacts according to the trait (i.e., how much the trait applies to the person). Names and traits varied between trials. The probe then asked if a certain name was at a certain position in the mental ranking (e.g., Is Sue second shiest?). For example, the participant should press “no” if Sue was the shiest person (see Figure 1). After the response or maximally 4 s, a 1-s break occurred, followed by a jittered inter-trial interval of 2 ± 0.5 s before the next trial started.

In the visual task, the stimuli consisted of 16 pictures of different fruits and vegetables, which naturally vary in color and size. Note that the stimuli were always presented in the same size, which was ensured by using the same amount of colored pixels to depict the objects against the grey background. The instruction was to sort the objects either with respect to color (from green to blue) or with respect to their natural size (from small to large). Both manipulation conditions appeared randomized and equally often. To ensure that participants rely on the same ranking of color and size, a uniform ranking was included in the preparation material (see procedure, Figure S1) and was rehearsed before the beginning of the EEG experiment.

In the verbal task, names were used as stimuli. The names were drawn randomly from a pool of 16 German names, which all differed in their first and last letter (Table S5). Participants should either sort the names in alphabetical order with respect to their first or last letter. Both manipulation conditions appeared randomized and equally often.

Each task and load condition was presented in a separate block of 16 task-specific trials. We also added 4 so-called order trials to each block to ensure that participants encoded all stimuli in the presented order and did not start to rank the stimuli before the manipulation period started (see Figure S2 for details). Importantly, the order trials were not included in any behavioral or EEG data analyses.

Each block was presented twice in randomized order during the whole EEG session, which (depending on the reaction time) lasted for 66.8 minutes (without breaks) maximum.

Thus, we recorded in total 240 trials, i.e., 32 task-specific trials per task and load condition. As our time window to analyze phase-amplitude coupling was 2.5 s long per trial, this number of trials leaves us with over 13 000 data points per analyzed phase segment, ensuring robust analyses (see *EEG data analyses and statistics/ Phase-amplitude coupling)*.

### Procedure and EEG recordings

After a successful screening, participants were invited to the EEG session. They were tested on the ranking of tasks (e.g., smallest to biggest object), which they had also received as preparation material (Table S3, Figure S1). They were provided with instructions and completed nine exercise trials for each of the tasks and load conditions.

Participants sat in a comfortable chair and performed the experimental task on the computer (Presentation ® software, version 20.1, build 12.04.17; NeuroBehavioral Systems), while EEG was recorded from 62 channels (Ag/AgCl scalp ring electrodes; Easycap ®) according to the 10-10 international system using a 64-channel amplifier (BrainAmp, Brain Products ®). Additionally, electrooculogram (EOG) was recorded from two channels placed at the left outer canthus and above the left eye to control for horizontal and vertical eye movements, respectively. The ground electrode was placed at Fpz. Depending on the Covid-19 restrictions, the data was referenced either to the tip of the nose or to both ear lobes. As all data was re-referenced to common average later in the preprocessing, the difference in reference should not be noticeable. The signal was recorded at a sampling rate of 1000 Hz and impedances were kept below 10 kΩ.

After the EEG recordings, participants filled out a survey (Table S6).

### Behavioural data analyses and statistics

The log files from Presentation ® were imported to Excel and descriptive data was extracted. Reaction times were calculated from the onset of the probe to the response (≤ 4 s) and included correct and incorrect responses. Accuracy in percentage was calculated by the number of correct responses divided by the total number of trials and multiplied with 100.

All 100 participants achieved a mean performance over all tasks significantly above chance level in the low (M = 89%, SD = 5.58) and high (M = 78%, SD = 6.63) load condition ^81^. However, some of the participants indicated after the EEG session that they reversed the given order of manipulation in a block (e.g., sorted the objects from big to small instead of from small to big in the visual task). Indeed, six participants reversed the order of a task as they performed significantly below chance level in at least one block (< 36% for the low and < 13% for the high load condition). Thus, these six participants had to be excluded from behavioral data analyses. As the order of the ranking does not affect brain activity, we did not exclude these participants from the EEG analyses.

For statistical analyses, we calculated repeated-measures ANOVAs for accuracy (in %) and reaction time (in ms) with the within-subject factors task x load and the between-subject factor autistic-traits group (N_low AQ_= 48, N_high AQ_ = 46) using IBM SPSS Statistics (version 29). Possible violations of the normal distribution should not have an effect on the robustness of the ANOVA ^82,83^. Sphericity was confirmed for reaction times, accuracy had to be Greenhouse-Geisser corrected. Post-hoc tests were performed based on estimated marginal means and if necessary, Bonferroni-corrected. We calculated post-hoc tests one-tailed if we had a directed hypothesis and two-tailed if we assumed no differences between the autistic-traits groups.

### EEG data analyses and statistics

#### EEG preprocessing

The continuous EEG data was preprocessed with BrainVision Analyzer 2.1 (Brain Products ®) to reduce and to control for artifacts as follows: In a visual inspection, large artifacts were removed from the data and if necessary, noisy channels were interpolated. Then the data was filtered between 0.1-100 Hz (48 dB/oct) and with a notch filter of 50 Hz. The EEG channels were re-referenced to a common average reference. Then an Independent Component Analysis for ocular correction was applied in order to remove horizontal (saccades) and vertical (blinks) eye movements. Additionally, noise on certain channels could be removed using ICA. On average 11 components (SD = 4.5) were removed per participant. Channels that were still too noisy for analyses after ICA were interpolated (in total, one channel was interpolated for 3 participants and 3 channels were interpolated for one participant). Last, the data was again visually inspected and all remaining artifacts were removed. Data of two individuals had to be excluded given that their data quality was insufficient. The continuous artifact-free data was then segmented in the 5-s manipulation periods (+ 250 ms in the beginning and in the end of the segment to counteract filter artifacts). The remaining 98 participants (49 participants per autistic-traits group) had on average between 29 and 31 manipulation period-trials (SD = 1.8-3.0) per task and load condition.

#### A priori source space transformation

These segments were then transformed into source space with the build-in module Low-Resolution Electromagnetic Tomography Analysis (LORETA) ^84,85^ in BrainVision Analyzer. Virtual electrodes were placed based on coordinates reported in neuroimaging studies for the social, verbal and visual working memory task ^19,21,22,49^: We defined the dorsomedial prefrontal cortex (DMPFC) as frontal region for FM-theta phase extraction. We defined 11 posterior regions of interest (ROIs) for gamma amplitude extraction: left and right temporal pole, left and right temporo-parietal junction, left and right inferior parietal lobe, left and right intraparietal sulcus, left, right and medial precuneus/posterior cingulate cortex (see Table S7). For each ROI, we defined all the voxels, that were found in relevant neuroimaging studies for the respective regions ^19,21,22,49^ (see Table S7). Then, the vectorial mean of the current density vectors across all the voxels included in one ROI was computed. X, y and z components of the mean current density vector were extracted from this value and averaged.

#### Power spectra

Only for the power spectra (but not for the phase-amplitude coupling analysis), the data was re-sampled to 512 Hz. The data was extracted from the first 2.5-s of the manipulation period. We only used the first 2.5 s of the 5-s manipulation period because participants indicated that they usually finished their mental ranking in the first half of the manipulation period in the low load condition (see Table S6). FFT power spectra (µV^2^) were computed from 1-256 Hz with a 1 Hz resolution using the full spectrum and a symmetric hanning window in BrainVision Analyzer.

#### Fitting oscillations & one over f

In order to determine if the FM-theta peak is a real oscillation, we divided the power spectrum into an aperiodic component (i.e., 1/f like characteristics) and in periodic compenents (i.e., peaks rising above the aperiod components) using the FOOOF toolbox ^50^ with a Matlab wrapper (Matlab R2016a) and Phyton version 3.5. For the frequency range of 1-15 Hz, we allowed an infinite number of peaks and defined the peak width between 2-12 Hz, the minimal peak height with 0.1 µV^2^ und peak threshold with 1.0 standard deviation of the aperiodic-removed power spectrum. We used a fixed aperiodic mode as our frequency range was rather small.

#### Complex Morlet wavelet transformation

7-cycle complex Morlet wavelet transformation was applied. For the instantaneous gamma amplitude, data was filtered between 30-70 Hz in 10 Hz steps. (30 Hz (26-34 Hz gauss), 40 Hz (34-46 Hz), 50 HZ (43-57 Hz), 60 (51-69 Hz), 70 (60-80 Hz)). As control analysis, also theta amplitude was extracted for the 4-7 Hz frequency range. For FM-theta phase, 5.5 Hz center frequency was filtered with a Morlet parameter of 3.8, thus resulting in a 4-7 Hz frequency band. These complex wavelet coefficients were transformed to absolute phase values (-pi to +pi) with the *complex data measures solution* module. After gamma amplitude and FM-theta phase extraction, single 5-s manipulation period-trials were exported from BrainVision Analyzer.

#### Statistical analyses of amplitudes

As control analysis, we calculated repeated-measures ANOVAs per task and frequency band for the amplitudes of the extracted (complex morlet wavelet transformation) frequency bands of the first 2.5 s of the 5-s manipulation period using IBM SPSS Statistics (version 29). For FM-theta, we used the amplitude from the DMPFC as dependent variable and the within-subject factor load and between-subject factor autistic-traits group. For the gamma amplitudes, we calculated the ANOVAs for the within-subject factors load and posterior ROIs and between-subject factor autistic-traits group. If the sphericity was violated, we used the Greenhouse-Geisser correction.

#### Phase-amplitude coupling

The data was further analyzed in Matlab (R2016a) using in-lab scripts ^27,86^. The posterior gamma amplitudes were z-transformed. The first 2.5-s of all manipulation period-trials were merged. As described above, we only used the first 2.5 s of the 5-s manipulation period because participants indicated that they usually finished their mental ranking in the first half of the manipulation period in the low load condition (see Table S6). This left us with roughly 375 data points per phase angle for averaging (i.e. roughly 5 theta cycles / s x 2.5 s manipulation period x 30 trials = 375 data points per phase angle, resulting in over 13 000 data points per analyzed phase segment). This number of data points ensures a robust analysis even with a relatively low number of trials.

Then, the z-transformed gamma amplitudes were sorted according to FM-theta phase angles (see Figure S4) and averaged into 10 theta phase segments in order to reduce data complexity. As the values were very small, we multiplied the data by 10 000 for analyses, so information was not lost due to rounding decimal places and a second time by 10 000 for the figures for better illustration purposes. For the circular plots, the data was transformed to percentage values in order to have the same scale across tasks and conditions.

#### Hierarchical generalized regression models

The z-transformed sorted gamma amplitudes were then further investigated by means of hierarchical generalized linear regression analysis conducted in R (version 4.2.2, R Studio 2022.07.1) ^87^. Hereby, possible violations of the normal distribution should have little impact on the robustness of regression coefficient estimation ^88^.

We analyzed regression models for each task and gamma frequency band separately with posterior gamma amplitude as criterion variable and FM-theta phase segments, load, *a priori* defined posterior ROIs, and autistic-traits groups as predictor variables with random intercepts for each individual. We were predominantly interested in models comprising any interaction between FM-theta phase segments and load, given that we hypothesized that FM-theta phase modulated posterior gamma amplitude depending on load.

The significance of an interaction was determined by comparing the second-order Aikaike information criterion (AICc) value of an interaction model to the AICc value of a model comprising only respective main effect terms. The smaller the AICc value of a model, the more variance of its criterion can be explained by its predictor variables. Thus, a positive difference in AICc values between an interaction and a main effect model indicates that the interaction model explains the criterion variable better than the main effect model. We only analyzed interaction models further that achieved an AICc >10 indicating a large effect ^89^ and if the corresponding main effect models were significant in comparison to a regression model without predictors and with random intercepts for each individual. We chose these conservative criteria in order to identify very strong and robust effects, i.e. to ensure that effects are true and not a product of testing multiple variables in complex interactions and separate regression models for tasks and frequency bands.

We found significant interactions between FM-theta phase segments, load condition and autistic-traits group (N = 49 participants per autistic-traits group) in specific gamma bands and tasks. As the predictor posterior ROIs did not reach significance in any combination with FM-theta phase, we averaged over posterior ROIs for further analyses.

#### Cosine-model fitting

For the significant regression models, we investigated whether there was a cosine relationship between the average posterior gamma amplitude and FM-theta phase segments as described in equation 1. Hereby, *Y* refers to individuals’ average posterior gamma amplitude, *X* to their FM-theta phase segment, α, β, γ, and δ to free cosine model parameter values denoting the cosine amplitude, frequency shift, phase shift and off-set, respectively. We set β to 1 to prevent an algorithmic solution neglecting our pre-defined FM-theta frequency. Then, α, γ, and δ parameter values were estimated using a Gauss-Newton fitting algorithm.

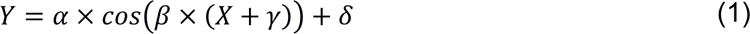

The model fit to the cosine models was determined by the mean absolute error (MAE). It describes the mean absolute deviation between predicted average posterior gamma amplitude values based on cosine model parameter estimates *ŷ* and individuals’ empirical average posterior gamma amplitude values *y* for each FM-theta phase segment. The smaller the error, the better the model fit.

Additionally, we compared the AICc value of the cosine model fit to an intercept model (i.e., the average over all data points – which should be around zero). While positive values already indicated that the cosine model tends to fit better, AICc > 2 indicate that the average posterior gamma amplitude estimates significantly fit to a cosine model and did not just randomly fluctuate around 0 ^89^. In this case, a significant effect (AICc > 2) was appropriate as criterion because we were not interested in multiple complex interaction effects but only tested whether the data fitted better to a cosine or an intercept model in pre-defined cases.

We determined the most plausible cosine phase-shifts between regression models of differential load and autistic-traits groups. Therefore, we computed the range of most plausible γ estimates by means of two-sided confidence intervals with a significance level of 5%. If 0° was not among the most plausible phase shift values, individuals’ gamma amplitude was likely shifted, i.e., significantly different from a null-shift reference model. If 0° was included in the confidence interval, the highest gamma amplitude was locked in the trough of FM-theta (i.e., 180°), i.e., resembled the null-shift reference model.

#### Circular-linear correlations

Circular-linear correlations between the FM-theta phase segment with the highest gamma amplitude (i.e., instantaneous FM-theta phase co-occurring with the maximum of sorted gamma amplitude) and AQ subscale scores over all participants as well as performance were computed ^86,90^. These correlations were only calculated for the tasks and frequency bands showing a significant phase-amplitude coupling.

## Acknowledgements

We thank Doris Schmid for support with programming and piloting the experimental design and Daniela Gresch for support with EEG data preprocessing. We are very grateful for the many valuable discussions with Anna Lena Biel and Carola Romberg-Taylor in order to design the experimental tasks. We give our special thanks to Jörg von Mankowski for providing tools to optimize our experimental design and advice during the complex analyses. We thank Eva Victoria Seegenschmiedt, Nele Habrecht and Ashley Yuan for assisting in designing the social questionnaire and the experimental tasks. This study was funded by a DFG grant to E.V.C.F. (FR 3961/1-1).

## Notes

### Competing Interest Statement

The authors have declared no competing interest.

### Summary of Updates

Revised manuscript includes additional analyses.

