## Supplemental Material for "Oscillatory brain activity as unified control mechanism for working memory and social cognition"

**Supplementary Material**

**Table S1. Participants.** Number and percentage of participants (Ps), mean, standard deviation (SD) and the range of the variables from the screening are indicated. **(A)** The low and high autistic-traits groups were defined by the Autism Spectrum Quotient (AQ; Baron-Cohen et al., 2001). Participants with a score ≤ 20 belong to the low autistic-traits group, whereas participants with scores > 20 belong to the high autistic-traits group. **(B)** The groups were comparable in terms of gender, age and education. **(C)** All participants from both groups were right-handed (short form of the Edinburgh Handedness Inventory; Oldfield, 1971), had normal or correct-to-normal (color) vision (Ishihara color test) and had German as mother tongue. Some of the participants were bi-lingual and had an additional mother tongue. **(D)** Ten out of the 100 participants reported that they have experienced anxiety within the last two years (equally distributed between autistic-traits groups). In the high autistic-traits group, one participant reported mild depression. We asked this person to complete the Beck Depression Inventory (BDI, version 1978; Beck et al., 1988) and this person scored 7 out of 63 points, which indicates no depression. Additionally, one person reported an Attention-Deficit Disorder but without medication. Three participants had a diagnosis of Autism Spectrum Disorder in the high autistic-traits group, however, their diagnosis did not lead to clinically significant impairment in daily-life functioning. No one took medication against psychiatric or neurological disorders, except for one person who used a low dosage of Escitalopram (7.5 mg). **(E)** Both groups achieved comparable scores in the verbal intelligence test (IQ) assessed with the Multiple Choice Vocabulary Test (MWT-B; Lehrl et al., 1995) and in visuospatial problem solving, assess with the Culture Fair Intelligence Test (CFT 20-R; Cattell and Cattell, 1960). **(F)** The German short version of the Big Five Inventory (BFI-K; Rammstedt and John, 2005) was applied to evaluate the Big Five personality traits. Compared to the high autistic-traits group, the low autistic-traits group showed a generally higher level in the Big Five personality traits ($N_{lowAQ} = 51$, $M_{lowAQ} = 3.69$, $SE_{lowAQ} = 0.06$ vs. $N_{highAQ} = 49$, $M_{highAQ} = 3.24$, $SE_{highAQ} = 0.06$, $F(1,98) = 29.17$, $p < 0.001$, partial $\eta^2 = 0.23$). These differences in personality traits are in accordance with a recent meta-analysis investigating the Big Five personality traits in Autism Spectrum Disorder (Lodi-Smith et al., 2019). Please note that emotional stability is the antonym of neuroticism, which means that all items were re-coded so that the direction of effects was consistent across Big Five traits.

| | Low Autistic-Traits Group | High Autistic-Traits Group |
|---|---|---|
| Number of Participants (Ps) | 51 | 49 |
| **(A) Autism Spectrum Quotient** | | |
| AQ: *mean (SD; range)* | 13.63 (4.24; 4-20) | 26.61 (5.21; 21-40) |
| **(B) Sociodemographics** | | |
| Gender: *number of Ps* | 25 female, 26 male | 27 female, 22 male |
| Age: *mean (SD; range) in years* | 23.61 (4.42; 19-39) | 23.55 (3.84; 18-34) |
| Education: *percentage of Ps* | 65% secondary school diploma, 35% university degree | 65% secondary school diploma, 35% university degree |
| **(C) Inclusion criteria** | | |
| *percentage of Ps:* | | |
| Handedness | 100% right-handed | 100% right-handed |
| Vision | 100% normal or corrected-to-normal | 100% normal or corrected-to-normal |
| Mother tongue | 100% German | 100% German |
| Additional second mother tongue | 16% | 8% |
| **(D) Medical history** | | |
| *number of Ps:* | | |
| Anxiety < 2 years ago | 5 | 5 |
| Mild depression < 2 years ago | 0 | 1 (BDI = 7: no depression) |
| Attention-Deficit Disorder | 0 | 1 |
| Autism-Spectrum Disorder (incl. Asperger; 1 had additionally mild anxiety 2 years ago) | 0 | 3 |
| Medication against psychiatric or neurological disorders | 0 | 1 (Escitalopram 7.5 mg) |
| **(E) IQ tests** | | |
| *mean (SD; range):* | | |
| MWT-B | 103.71 (11.49; 88-145) | 106.08 (9.83; 93-136) |
| CFT 20-R | 114.27 (13.17; 86-145) | 118.45 (13.18; 90-142) |
| **(F) Big 5 personality traits** | | |
| *mean (SD; range):* | | |
| Extraversion | 3.73 (0.77; 1.75-4.75) | 2.89 (0.92; 1.25-4.25) |
| Agreeableness | 3.27 (0.78; 1.50-4.75) | 2.85 (0.78; 1.00-4.25) |
| Conscientiousness | 3.84 (0.69; 1.50-5.00) | 3.56 (0.72; 1.75-5.00) |
| Emotional Stability | 3.35 (0.82; 1.50-5.00) | 2.93 (0.95; 1.25-4.75) |
| Openness to Experience | 4.21 (0.63; 2.40-5.00) | 3.96 (0.70; 2.60-5.00) |

**Table S2. Selection of Traits.** We selected 99 adjectives based on Grühn and Smith (2008) that described a trait and had to be inferred from the person's behavior. We excluded all items that referred to a state (such as angry), a description (such as young) or a direct behavior (such as active). We furthermore only considered adjectives that were rated > 1.5 valence points on the scale of 1-7 in the list from Grühn and Smith (2008) as our participants should rate friends and family. So, items like brutal and cruel for example were excluded. We used German traits with 4-12 characters. We then tested these 99 traits in pretests and selected the final 48 traits based on how well they described different aspects of personality. The final 48 traits are as follows:

| Number of Trait | German | English |
|---|---|---|
| **1** | analytisch | analytical |
| **2** | angepasst | conformist |
| **3** | arrogant | arrogant |
| **4** | aufbrausend | quick-tempered |
| **5** | autoritär | authoritarian |
| **6** | beharrlich | insistent |
| **7** | berechnend | calculating |
| **8** | besonnen | level-headed |
| **9** | dominant | dominant |
| **10** | ehrgeizig | ambitious |
| **11** | ehrlich | honest |
| **12** | eigenwillig | willful |
| **13** | einfallslos | unimaginative |
| **14** | einfühlsam | empathic |
| **15** | eitel | vain |
| **16** | emotional | emotional |
| **17** | empfindlich | touchy |
| **18** | entschlossen | determined |
| **19** | flexibel | flexible |
| **20** | friedlich | peaceful |
| **21** | furchtsam | frightened |
| **22** | gehemmt | inhibited |
| **23** | gerecht | just |
| **24** | gesellig | sociable |

| Number of Trait | German | English |
|---|---|---|
| **25** | gütig | kind |
| **26** | höflich | polite |
| **27** | impulsiv | impulsive |
| **28** | intelligent | intelligent |
| **29** | ironisch | ironic |
| **30** | kämpferisch | belligerent |
| **31** | kritisch | critical |
| **32** | launisch | moody |
| **33** | liberal | liberal |
| **34** | rational | rational |
| **35** | reif | mature |
| **36** | reizbar | irritable |
| **37** | sachlich | factual |
| **38** | schüchtern | shy |
| **39** | sensibel | sensitive |
| **40** | sentimental | sentimental |
| **41** | spontan | spontaneous |
| **42** | stolz | proud |
| **43** | stur | stubborn |
| **44** | taktvoll | tactful |
| **45** | treu | loyal |
| **46** | unsicher | unsure |
| **47** | verspielt | prettily |
| **48** | zaghaft | timid |

**Table S3. Social Questionnaire for the Social Task in the EEG Experiment.** In the screening, participants filled out an individualized questionnaire (i.e., social questionnaire). We used this questionnaire to build individualized trials for the social task in the EEG experiment. We programmed the social questionnaire on the platform *soscisurvey.de*. In the social questionnaire, we first asked participants to name 10 contact persons such as close friends and family members. We then displayed each trait with a short description and asked the participants to rate their 10 contact persons on how much this trait applied to each contact person from 0 to 100% in 5% steps. The low autistic-traits group needed on average 39 minutes to fill out the social questionnaire, whereas it took the high autistic-traits group 51 minutes. The social questionnaire was completed about two weeks prior to the EEG recordings in both groups. Before the EEG recordings, we asked participants to review the description of the traits again in the preparation material - which was sent two days prior to the EEG recordings - and during the instructions. Thus, participants had the same criteria in mind to rank their contact persons. For the social task in the EEG experiment, participants were asked to infer the traits from their contact persons to mentally rank the names according to the traits.

| | **Low Autistic-Traits Group** | **High Autistic-Traits Group** |
|---|---|---|
| **Social questionnaire** *mean (SD; range):* | | |
| Processing time *(in minutes)* | 38.59 (8.53; 22-56) | 50.57 (26.08; 26-192) |
| Time between the social questionnaire and the EEG recording *(in days)* | 14.78 (5.76; 6-27) | 13.06 (4.31; 7-25) |

**Table S4. Social Questionnaire Re-test Reliability.** We evaluated the re-test reliability of our social questionnaire (i.e., individualized questionnaire of the screening to build the trials for the social task in the EEG experiment) in a separate study. We asked 63 participants to fill out the social questionnaire online twice, evaluating the same contact persons in both sessions. Four participants had to be excluded. The final sample included 35 participants with low autistic-traits based on the Autism Spectrum Quotient (AQ; Baron-Cohen et al., 2001: M = 13.74, SD = 3.73). The low autistic-traits group included 30 female and 5 male participants who were on average 22.34 years old (SD = 4.24) and had 14.89 days (SD = 4.53) between the two sessions. It also included 24 participants with high autistic-traits (AQ: M = 27.08, SD = 6.06). The high autistic-traits group included 18 female, 4 male and 2 divers participants who were on average 22.88 years old (SD = 4.70) and had 15.75 days (SD = 4.22) between the two sessions.

The data from the social questionnaire was analyzed in the same way as we did for the social task for the EEG recordings: We only included traits for which participants had ranked their contacts ≥ 15 points apart from one another. Then, we calculated repeated-measures correlation (repeated measures being the 10 contact persons) separately for every trait with > 15 participants in each autistic-traits group between sessions 1 and 2 in R (version 4.2.2, R Studio 2022.12.0; R Core Team, 2022). The value of the repeated measures correlation coefficient (r), the 95% confidence interval (CI) for the repeated measures coefficient and the actual number of participants are displayed for all 32 traits, which fulfilled our criteria (i.e., traits for which at least 15 participants have ranked their contact persons ≥ 15 points apart from one another).

The low autistic-traits group reached a mean re-test reliability of r = 0.75 (range between 0.61 and 0.86). The high autistic-traits group reached a mean re-test reliability of r = 0.73 (range between 0.59 and 0.85). All confidence intervals between the low and high autistic-traits groups overlap, except for the one of trait number 35 (“reif”/“mature”), indicated with an *.

| Number of Trait | Trait (German) | Low Autistic-Traits Group | | | High Autistic-Traits Group | | |
|---|---|---|---|---|---|---|---|
| | | r | 95% CI | N | r | 95% CI | N |
| 1 | analytisch | 0.811 | 0.766 - 0.848 | 30 | 0.746 | 0.674 - 0.805 | 20 |
| 4 | aufbrausend | 0.795 | 0.745 - 0.836 | 28 | 0.764 | 0.696 - 0.819 | 20 |
| 6 | beharrlich | 0.697 | 0.614 - 0.765 | 20 | 0.646 | 0.550 - 0.726 | 19 |
| 7 | berechnend | 0.703 | 0.631 - 0.764 | 25 | 0.664 | 0.562 - 0.746 | 16 |
| 9 | dominant | 0.837 | 0.792 - 0.873 | 24 | 0.818 | 0.761 - 0.862 | 19 |
| 12 | eigenwillig | 0.775 | 0.708 - 0.829 | 19 | 0.632 | 0.523 - 0.721 | 16 |
| 14 | einfühlsam | 0.766 | 0.703 - 0.817 | 23 | 0.809 | 0.750 - 0.855 | 19 |
| 15 | eitel | 0.764 | 0.698 - 0.818 | 21 | 0.729 | 0.655 - 0.790 | 21 |
| 17 | empfindlich | 0.783 | 0.727 - 0.829 | 25 | 0.797 | 0.731 - 0.848 | 17 |
| 18 | entschlossen | 0.733 | 0.665 - 0.789 | 24 | 0.627 | 0.520 - 0.714 | 17 |
| 19 | flexibel | 0.614 | 0.524 - 0.691 | 24 | 0.725 | 0.637 - 0.794 | 16 |
| 20 | friedlich | 0.816 | 0.765 - 0.857 | 23 | 0.779 | 0.711 - 0.833 | 18 |
| 21 | furchtsam | 0.796 | 0.741 - 0.840 | 24 | 0.765 | 0.688 - 0.825 | 16 |
| 22 | gehemmt | 0.768 | 0.707 - 0.817 | 24 | 0.742 | 0.661 - 0.805 | 17 |
| 24 | gesellig | 0.822 | 0.771 - 0.862 | 22 | 0.827 | 0.773 - 0.869 | 19 |
| 25 | gütig | 0.669 | 0.582 - 0.741 | 21 | 0.788 | 0.718 - 0.843 | 16 |
| 27 | impulsiv | 0.627 | 0.532 - 0.706 | 21 | 0.724 | 0.642 - 0.790 | 18 |

| | | | | | | | |
|---|---|---|---|---|---|---|---|
| 30 | kämpferisch | 0.749 | 0.676 - 0.807 | 20 | 0.693 | 0.598 - 0.769 | 16 |
| 31 | kritisch | 0.651 | 0.569 - 0.720 | 25 | 0.594 | 0.484 - 0.685 | 18 |
| 32 | launisch | 0.783 | 0.729 - 0.827 | 27 | 0.796 | 0.732 - 0.847 | 18 |
| 34 | rational | 0.711 | 0.641 - 0.769 | 26 | 0.724 | 0.642 - 0.790 | 18 |
| 35* | reif | 0.861 | 0.816 - 0.895 | 19 | 0.741 | 0.658 - 0.807 | 16 |
| 36 | reizbar | 0.794 | 0.742 - 0.836 | 27 | 0.778 | 0.711 - 0.831 | 19 |
| 37 | sachlich | 0.751 | 0.685 - 0.805 | 23 | 0.761 | 0.686 - 0.821 | 17 |
| 38 | schüchtern | 0.756 | 0.692 - 0.808 | 24 | 0.849 | 0.798 - 0.888 | 17 |
| 39 | sensibel | 0.652 | 0.562 - 0.727 | 21 | 0.699 | 0.618 - 0.765 | 21 |
| 40 | sentimental | 0.648 | 0.561 - 0.720 | 23 | 0.666 | 0.574 - 0.742 | 19 |
| 41 | spontan | 0.739 | 0.677 - 0.790 | 28 | 0.68 | 0.584 - 0.756 | 17 |
| 43 | stur | 0.803 | 0.748 - 0.848 | 22 | 0.727 | 0.648 - 0.791 | 19 |
| 46 | unsicher | 0.807 | 0.756 - 0.848 | 25 | 0.728 | 0.641 - 0.797 | 16 |
| 47 | verspielt | 0.707 | 0.638 - 0.764 | 27 | 0.707 | 0.624 - 0.775 | 19 |
| 48 | zaghaft | 0.737 | 0.679 - 0.785 | 32 | 0.618 | 0.506 - 0.709 | 16 |

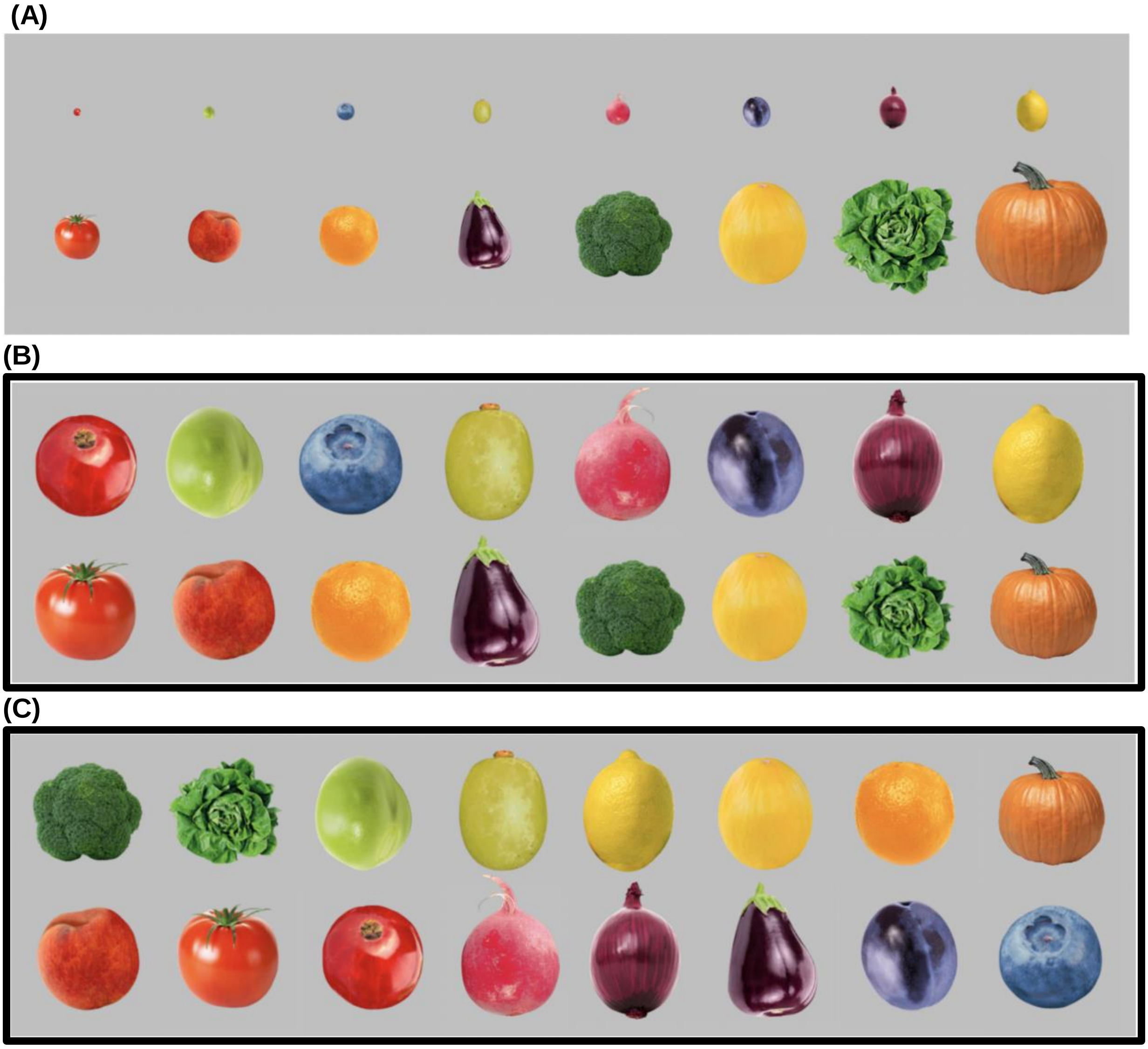


**Figure S1. Visual Stimuli. (A)** We used 16 pictures of different fruits and vegetables that naturally vary in size and color. In the pretests, we tested various designs for the visual task and found comparable performance for the subtasks “rank the objects according to size from small to big” and “according to color from green to blue”. Importantly, the subtasks were only comparable when displaying all objects in the same size and not in their actual size. The final stimuli (indicated by a black frame) were thus all presented in the same size - which was ensured by using the same amount of colored pixels to depict the objects against the grey background - as illustrated in (B) and (C): Objects sorted according to **(B)** size from small to big and **(C)** color from green to blue. The objects and their sorting instruction varied between trials. The ranking of the fruits and vegetables were sent to the participants in the preparation material two days prior to the EEG recording. Before the EEG recording, participants could review the ranking in the instruction and had to sort these objects themselves in a practical test. This ensured that the participants had the same sorting-criteria in mind. Additionally, participants were instructed to imagine the objects visually such as a photograph in the visual task and infer color and size from this mental picture.

**Table S5. Verbal Stimuli.** We used 16 different German first names, which all differed in their first and last letter. They were half female and half male names, consisted of 4-6 letters and had no ambiguity how to spell them in respect to their first and last letter. Names and their sorting instruction varied between trials. Before the EEG recording, participants were asked to write down the alphabet to ensure that they were able to alphabetically sort the names by their first and last letter. Additionally, participants were instructed to encode and manipulate the verbal stimuli verbally in their mind.

| | | | |
|---|---|---|---|
| Astrid | Felix | Jakob | Rolf |
| Birgit | Gustav | Karin | Sabine |
| Daniel | Herwig | Leni | Theo |
| Esther | Iris | Moritz | Ursula |

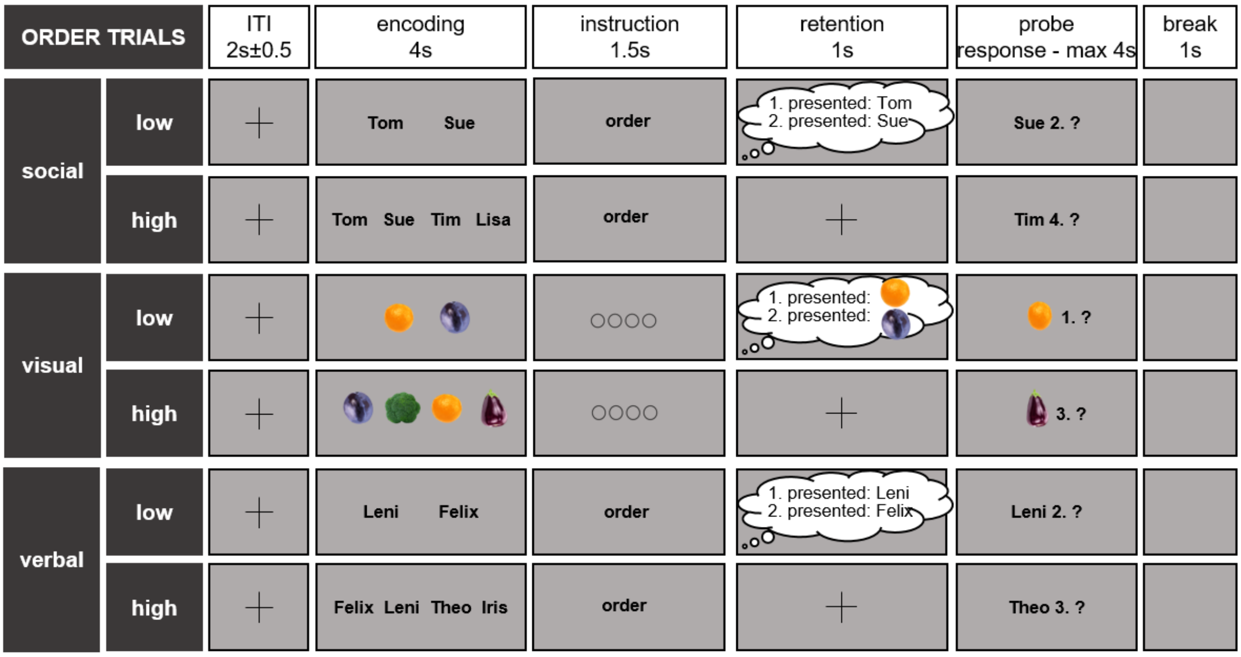


**Figure S2. Order Trials.** We added so-called order trials to each block of task-specific trials (i.e., trials in the social, visual and verbal task, see Figure 1 in the main manuscript for the task-specific trials). As in the task-specific trials, either two or four stimuli were presented for 4 s for the participants to encode. The encoding stimuli were the same as for the task-specific trials (e.g., names as stimuli for the social and verbal tasks or pictures of fruits and vegetables for the visual task). However, instead of the task-specific instruction, participants were asked to remember the stimuli in the presented order without any manipulation. After the 1-s retention period (instead of a 5-s manipulation period as in the task-specific trials), the probe then referred to the original position of the stimuli during the encoding period. The participants pressed yes or no with their right hand as fast and accurately as possible on the left and right mouse key, respectively. Match and non-match probes were randomized and occurred equally often. After the response or maximally after 4 s, there was a 1-s break before the next trial started with an inter-trial interval (ITI) jittered around 2 s (± 0.5 s). The order trials should ensure that participants encoded all stimuli in the presented order and did not start to rank the stimuli before the manipulation period started. Importantly, the order trials were not included in any behavioral or EEG data analyses.

**Table S6. Post-EEG-Recording Survey.** The 100 participants filled out a survey after the EEG recording. Data is missing from one person from the low autistic-traits group, thus we have data from 50 low autistic-traits and 49 high autistic-traits participants. Among other questions, we asked the participants how much time they actually needed to rank the stimuli in their head during the manipulation period. Over 50% of the participants in both groups indicated that they finished the ranking in their mind during the 5-s manipulation period already about 2 s before the probe appeared in the low load condition. This is the reason why we only analyzed the first half of the manipulation period.

| | **Low Autistic-Traits Group** | **High Autistic-Traits Group** |
|---|---|---|
| **Manipulation period - timing:**<br>*(percentage of participants indicating the specified time class)* | | |
| In the low load condition, the ranking of stimuli in my mind was completed… | 19% about 3 s before the probe<br>38% about 2 s before the probe<br>19% about 1 s before the probe<br>9% shortly before the probe<br>10% after the probe<br>0% never<br>5% other | 14.3% about 3 s before the probe<br>40% about 2 s before the probe<br>14.3% about 1 s before the probe<br>14.3% shortly before the probe<br>13% after the probe<br>0% never<br>4% other |
| In the high load condition, the ranking of stimuli in my mind was completed… | 0% about 3 s before the probe<br>5% about 2 s before the probe<br>13% about 1 s before the probe<br>30% shortly before the probe<br>44% after the probe<br>4 % never<br>4% other | 0% about 3 s before the probe<br>2% about 2 s before the probe<br>14% about 1 s before the probe<br>34% shortly before the probe<br>42% after the probe<br>7% never<br>1% other |

**Table S7. Regions of Interest (ROIs).** The x, y, z coordinates are indicated according to the Montreal Neurological Institute (MNI). We based the coordinates on Meyer et al. (2015, 2012), Meyer and Collier (2020) and Todd and Marois (2004). If the indicated literature reported the coordinates in Talairach coordinates, we transferred them to MNI with SLORETA software (Standardized Low Resolution Electromagnetic Tomography; sLORETA v20190617; Pascual-Marqui, 2002; Pascual-Marqui, 2007).

**(A)** We defined the dorsomedial prefrontal cortex (DMPFC) as frontal ROI, from which we extracted the FM-theta phase. **(B)** We defined 11 posterior ROIs, from which we extracted gamma amplitude values. The ROIs written in purple were associated with social working memory in the cited studies. The ROIs written in blue were found to be active in non-social tasks or responsible for general load effects in working memory processes.

| ROI name | ROI abbreviation | X (MNI) | Y (MNI) | Z (MNI) | Reference |
|---|---|---|---|---|---|
| **(A)** | | | | | |
| dorsomedial prefrontal cortex | DMPFC | 12 | 29 | 31 | Meyer et al. (2012) |
| | | -12 | 38 | 49 | Meyer et al. (2012) |
| | | 15 | 39 | 54 | Meyer et al. (2015) |
| | | 6 | 54 | 24 | Meyer et al. (2015) |
| | | 12 | 66 | 15 | Meyer et al. (2015) |
| | | -9 | 54 | 39 | Meyer et al. (2015) |
| | | 12 | 36 | 54 | Meyer and Collier (2020) |
| | | -6 | 34 | 56 | Meyer and Collier (2020) |
| | | -4 | 44 | 48 | Meyer and Collier (2020) |
| | | -8 | 54 | 36 | Meyer and Collier (2020) |
| | | 14 | 38 | 54 | Meyer and Collier (2020) |
| | | -10 | 58 | 26 | Meyer and Collier (2020) |
| **(B)** | | | | | |
| left temporal pole | lTP | -50 | 10 | -32 | Meyer and Collier (2020) |
| | | -62 | -10 | -22 | Meyer and Collier (2020) |
| | | -60 | -2 | -24 | Meyer and Collier (2020) |
| right temporal pole | rTP | 48 | 12 | -36 | Meyer and Collier (2020) |
| | | 52 | 16 | -28 | Meyer and Collier (2020) |
| | | 62 | -6 | -18 | Meyer and Collier (2020) |
| left temporo-parietal junction | lTPJ | -42 | -70 | 40 | Meyer et al. (2012) |
| | | -52 | -64 | 40 | Meyer and Collier (2020) |
| | | -48 | -58 | 30 | Meyer and Collier (2020) |
| | | -56 | -64 | 26 | Meyer and Collier (2020) |
| | | -54 | -66 | 24 | Meyer and Collier (2020) |
| | | -42 | -62 | 24 | Meyer and Collier (2020) |
| | | -50 | -66 | 16 | Meyer and Collier (2020) |
| right temporo-parietal junction | rTPJ | 42 | -54 | 24 | Meyer et al. (2015) |
| | | 54 | -66 | 27 | Meyer et al. (2015) |
| | | 52 | -60 | 44 | Meyer and Collier (2020) |

| | | | | | |
|---|---|---|---|---|---|
| left inferior parietal lobe | lIPL | -33 | -54 | 45 | Meyer et al. (2015) |
| right inferior parietal lobe | rIPL | 45 | -33 | 45 | Meyer et al. (2015) |
| | | 33 | -66 | 48 | Meyer et al. (2015) |
| | | 44 | -36 | 44 | Meyer and Collier (2020) |
| left intraparietal sulcus | lIPS | -22 | -69 | 42 | Todd and Marois (2004) |
| right intraparietal sulcus | rIPS | 23 | -63 | 46 | Todd and Marois (2004) |
| medial precuneus/ posterior cingulate cortex | PC/PCC | 0 | -61 | 46 | Meyer et al. (2012) |
| | | -3 | -60 | 27 | Meyer et al. (2015) |
| | | -3 | -54 | 21 | Meyer et al. (2015) |
| | | 3 | -54 | 36 | Meyer et al. (2015) |
| | | -2 | -50 | 28 | Meyer and Collier (2020) |
| | | 0 | -68 | 36 | Meyer and Collier (2020) |
| left precuneus/ posterior cingulate | lPC/PCC | -10 | -60 | 54 | Meyer and Collier (2020) |
| right precuneus/ posterior cingulate | rPC/PCC | 12 | -62 | 58 | Meyer and Collier (2020) |
| | | 4 | -42 | 46 | Meyer and Collier (2020) |

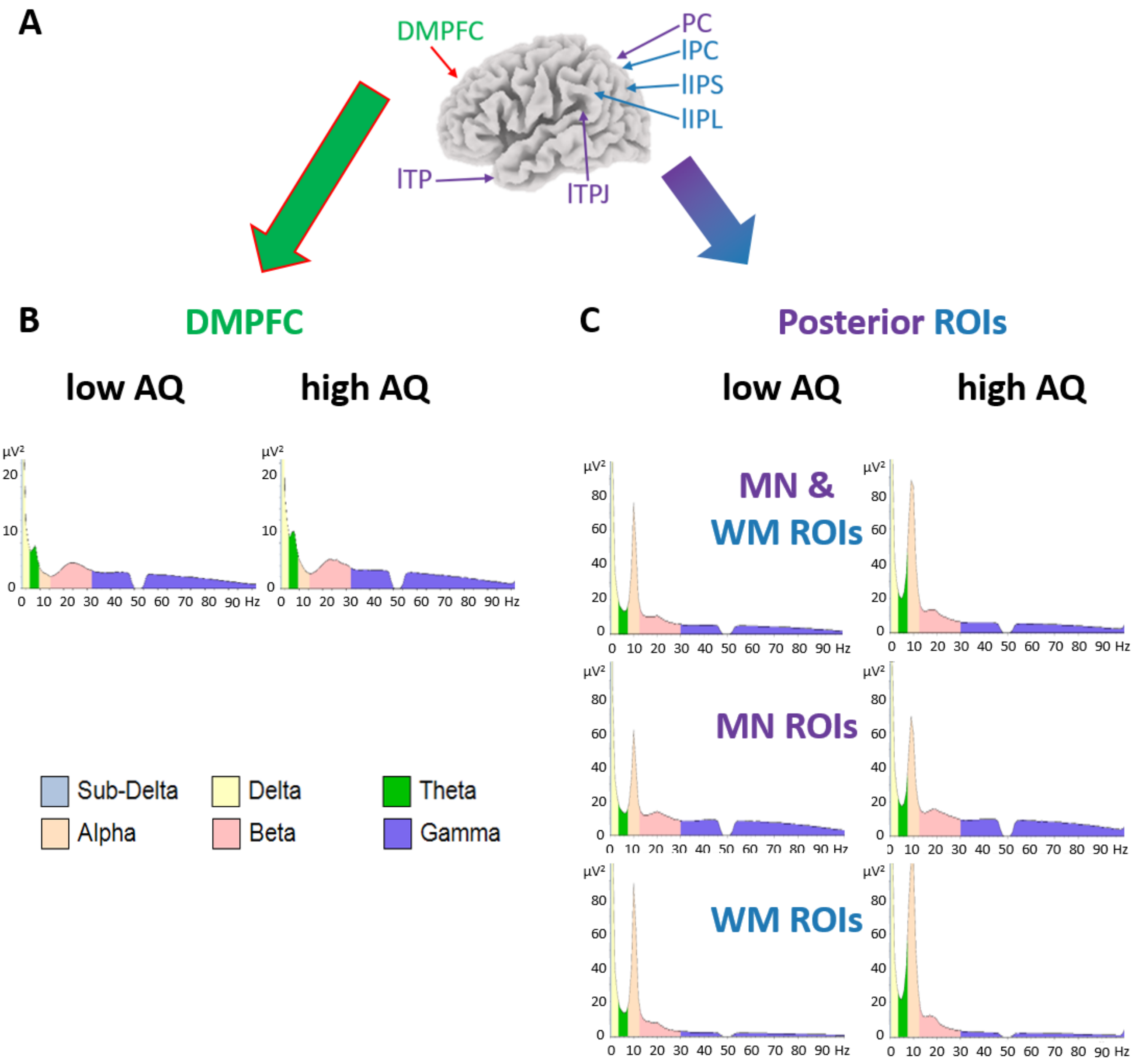


**Figure S3. Power Spectra. (A)** Data was extracted from one frontal ROI (i.e., dorsomedial prefrontal cortex (DMPFC, green/red) and eleven posterior ROIs. Five of these posterior ROIs were reported to be active during social working memory tasks (left/right temporal pole (l/rTP), left/right temporo-parietal junction (l/rTPJ), medial precuneus (PC)) and match with regions from the mentalizing network (MN, in purple; Meyer et al., 2015, 2012; Meyer and Collier, 2020). Six of these regions were found to be active in nonsocial tasks or responsible for general load effects in working memory processes (left/right precuneus/posterior cingulate cortex (l/rPC), left/right inferior parietal lobe (l/rIPL), left/right intraparietal sulcus (l/rIPS)) and considered typical working memory regions (WM, in blue; Meyer et al., 2015; Meyer and Collier, 2020; Todd and Marois, 2004). The arrows show the approximate left and medial ROIs, for all coordinates see Table S7.

**(B)** Power (µV$^2$) averaged over the first 2.5-s of the manipulation period (y- axis) are shown for the freuqency range of 1-100 Hz (x-axis) with a 1 Hz resolution for the DMPFC, seperatley for the low (left column) and high (right colum) autistic-traits (AQ) groups. A clear theta peak (green) is visible in the power spectra at the DMPFC for both groups.

**(C)** Power ($\mu V^2$) averaged over the first 2.5-s of the manipulation period (y- axis) are shown for the freuqency range of 1-100 Hz (x-axis) with a 1 Hz resolution, seperatley for the low (left column) and high (right colum) autistic-traits (AQ) groups. In the first row, power was averaged over all eleven posterior ROIs (MN & WM ROIs). In the second row, power was averaged over the five posterior ROIs associated with the mentalizing network (MN ROIs). In the third row, power was averaged over the six posterior ROIs associated with the working memory system (WM ROIs). No theta peaks are visible in the power spectra at the posterior ROIs.

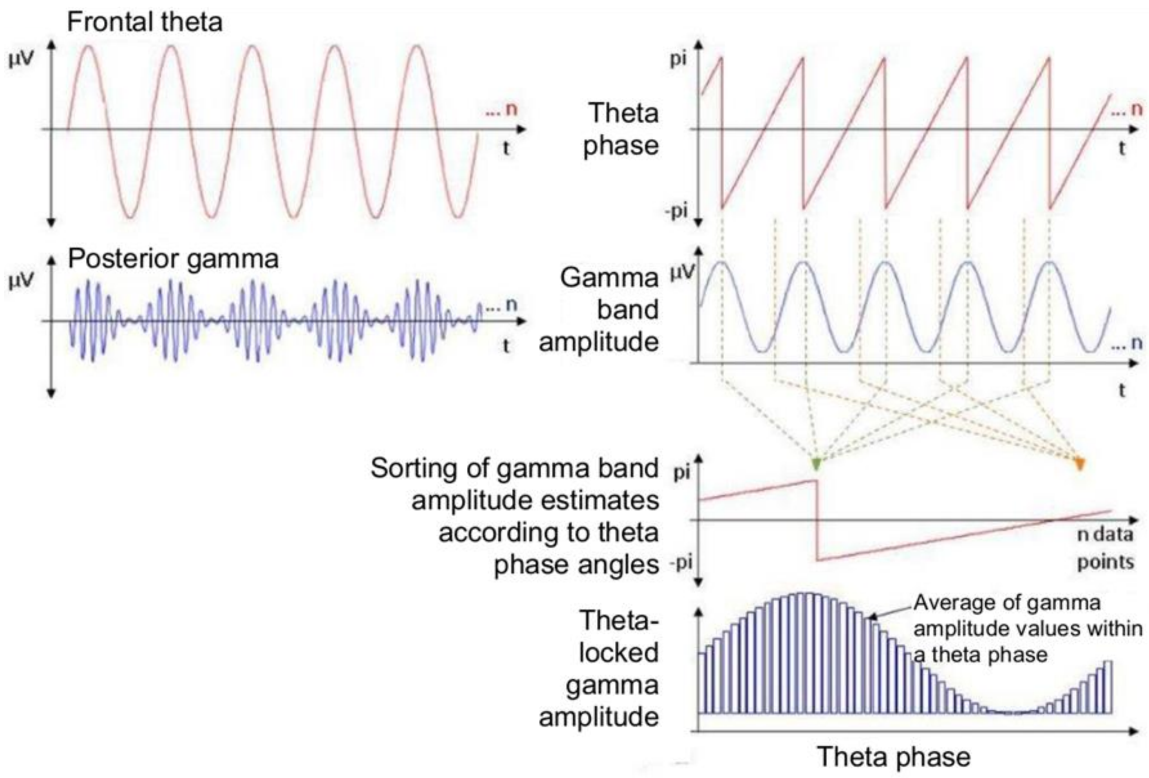


**Figure S4. Analysis for Phase-Amplitude Coupling.** Using complex Morlet wavelet transformation, FM-theta absolute phase values (-pi to +pi) were extracted from the dorsomedial prefrontal cortex (DMPFC) and instantaneous gamma band amplitude values were extracted from 11 posterior ROIs (see Table S7). Then the posterior gamma amplitudes were z-transformed and sorted according to the FM-theta phase angles. This resulted in averaged posterior gamma amplitude values related to different FM-theta phase angles.

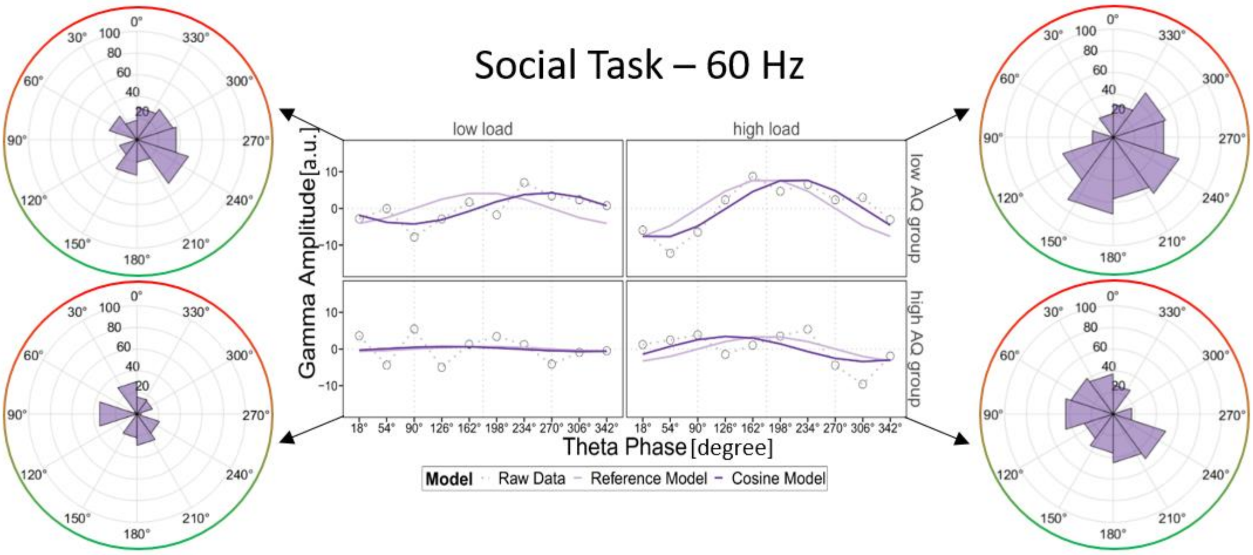


**Figure S5. DMPFC Phase-Amplitude Coupling in the Social Task.** FM-theta phase was extracted from the dorsomedial prefrontal cortex (DMPFC). 60-Hz posterior gamma amplitude was extracted from 11 posterior regions of interest (ROIs, see Table S7). The z-transformed posterior instantaneous gamma amplitude was sorted according to instantaneous FM-theta phase and averaged over all 11 posterior ROIs. In the line charts, the grey dots indicate the empirical z-transformed and sorted 60-Hz gamma amplitudes, the light purple lines our null-shift reference cosine model (simulating that strongest gamma amplitudes were locked in the trough of FM-theta phase) and the dark purple lines the cosine model fitted to our empirical data. In the circular plots, z-transformed sorted gamma amplitudes (purple) are displayed as percentage of the signal, FM-theta peak (0°) is indicated in red and FM-theta trough (180°) in green.
In the social task in the here displayed 60-Hz frequency band, the phase-amplitude coupling was similar but slightly weaker than in the 70-Hz frequency band, described in the main manuscript and illustrated in Figure 3C.

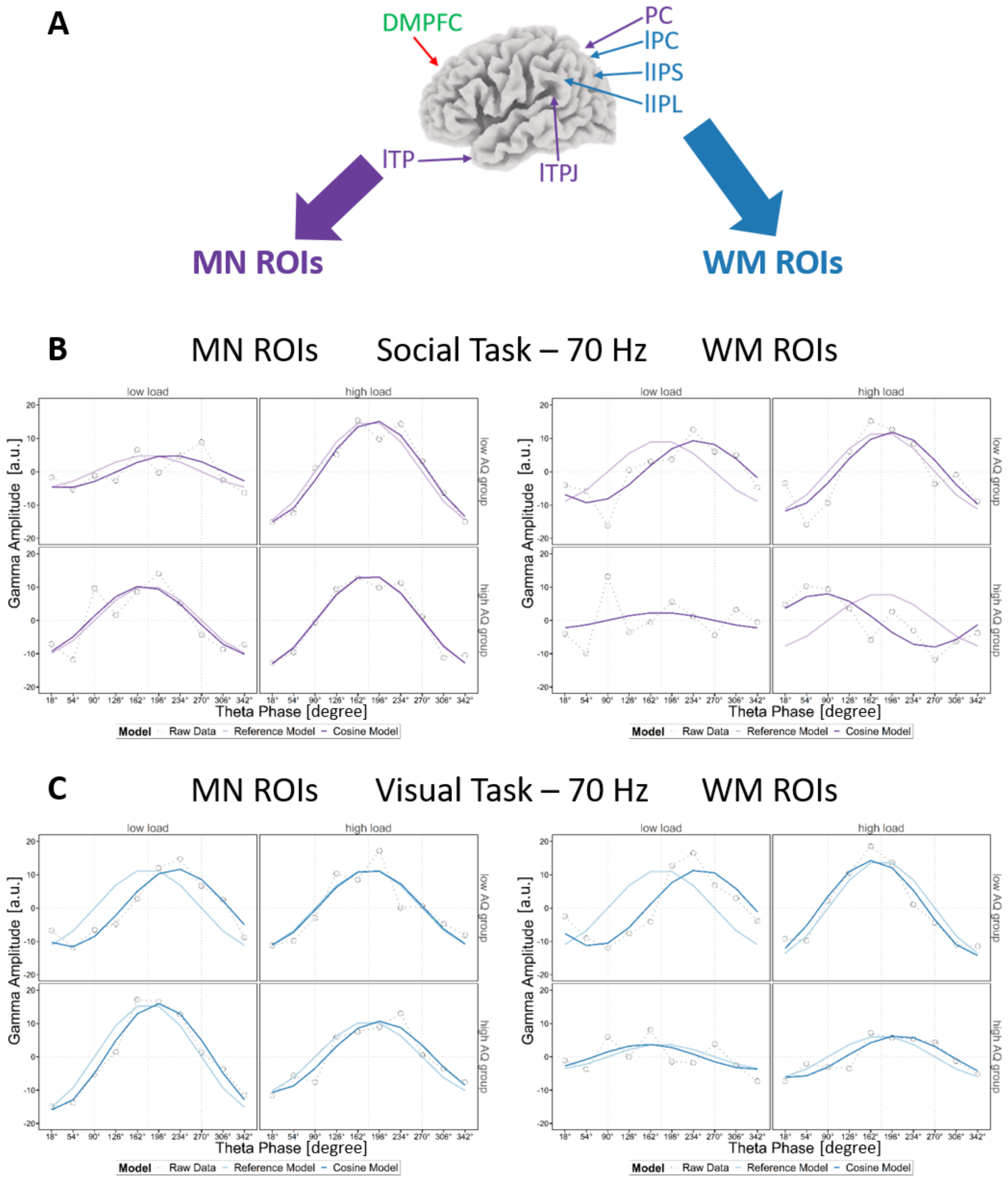


**Figure S6. DMPFC Phase-Amplitude Coupling separately for posterior ROIs associated with the mentalizing network (MN) and working memory system (WM).**

**(A)** FM-theta phase was extracted from the dorsomedial prefrontal cortex (DMPFC; green/red). 70-Hz posterior gamma amplitude was extracted from 11 posterior regions of interest. Five of these posterior ROIs were reported to be active during social working memory tasks (left/right temporal pole (l/rTP), left/right temporo-parietal junction (l/rTPJ), medial precuneus (PC)) and match with regions from the mentalizing network (MN, in purple; Meyer et al., 2015, 2012; Meyer and Collier, 2020). Six of these regions were found to be active in non-social tasks or responsible for general load effects in working memory processes (left/right precuneus/posterior cingulate cortex (l/rPC), left/right inferior parietal lobe (l/rIPL), left/right intraparietal sulcus (l/rIPS)) and considered typical working memory regions (WM, in blue; Meyer et al., 2015; Meyer and Collier, 2020; Todd and Marois, 2004). The arrows show the approximate left and medial ROIs, for all coordinates see Table S7.

**(B)** and **(C)** The z-transformed posterior instantaneous gamma amplitude was sorted according to instantaneous FM-theta phase and averaged either over the ROIs associated with the mentalizing network (left) or the working memory system (right).
In the single line charts, the grey dots indicate the empirical z-transformed and sorted 70-Hz gamma amplitudes, the light purple (i.e., social task (B)) or blue (i.e., visual task (C)) lines our null-shift reference cosine model (simulating that strongest gamma amplitudes were locked in the trough of FM-theta phase) and the dark purple (i.e., social task (B)) or blue (i.e., visual task (C)) lines the cosine model fitted to our empirical data.
In the social (B) and visual (C) tasks, in the low autistic-traits (AQ) group (top rows), the same pattern emerged in the phase-amplitude coupling with mentalizing and working memory regions: In the low load condition, strongest gamma amplitude was shifted towards 270°, while the maximal gamma amplitude was locked to the FM-theta phase in the high load condition.
In the high autistic-traits (AQ) group (bottom rows), the load-dependent phase-amplitude coupling (as obtained for individuals with low autistic traits), could not be systematically found.

**Table S8. Statistical Amplitude Analysis. (A)** We calculated repeated-measures ANOVA for FM-theta amplitude at the DMPFC with the within-subject factor load and between-subject factor autistic-traits (AQ) group separately for the tasks. As expected, FM-theta amplitude was higher in the high than low load condition in all tasks. Importantly, there was no difference in FM-theta amplitude between the autistic-traits groups and no significant interaction in any task. **(B)** We calculated repeated-measures ANOVA for gamma amplitude separately for the tasks and the center frequencies of 60 Hz and 70 Hz with the within-subject factor load and posterior ROIs and the between-subject factor autistic-traits (AQ) group. There was no main effect in load or autistic-traits groups and no significant interaction between these factors or involving posterior ROIs in any task. The main effect ROIs became significant in all tasks and frequency bands, however, this effect is trivial and to be expected. Importantly, it was not systematically modulated by load or autistic-traits group. Moreover, the z-transformation of gamma amplitudes for the phase-amplitude coupling eliminated this effect.

| **(A) FM-theta (4-7 Hz) amplitude at DMPFC** | | |
|---|---|---|
| **4-7 Hz** | | |
| verbal | load | $F(1, 96) = 16.418$, $p < 0.001$, partial $\eta^2 = 0.146$ |
| | AQ group | $F(1, 96) = 2.386$, $p = 0.126$, partial $\eta^2 = 0.024$ |
| | load x AQ group | $F(1, 96) = 0.043$, $p = 0.836$, partial $\eta^2 = 0.000$ |
| social | load | $F(1, 96) = 4.808$, $p = 0.031$, partial $\eta^2 = 0.048$ |
| | AQ group | $F(1, 96) = 3.506$, $p = 0.064$, partial $\eta^2 = 0.035$ |
| | load x AQ group | $F(1, 96) = 0.260$, $p = 0.611$, partial $\eta^2 = 0.003$ |
| visual | load | $F(1, 96) = 24.934$, $p < 0.001$, partial $\eta^2 = 0.206$ |
| | AQ group | $F(1, 96) = 2.616$, $p = 0.109$, partial $\eta^2 = 0.027$ |
| | load x AQ group | $F(1, 96) = 0.054$, $p = 0.817$, partial $\eta^2 = 0.001$ |

| **(B) Gamma amplitude at posterior ROIs** | | |
|---|---|---|
| **60 Hz** | | |
| social | load | $F(1, 96) = 0.659$, $p = 0.419$, partial $\eta^2 = 0.007$ |
| | AQ group | $F(1, 96) = 0.611$, $p = 0.436$, partial $\eta^2 = 0.006$ |
| | load x AQ group | $F(1, 96) = 0.003$, $p = 0.953$, partial $\eta^2 = 0.000$ |
| | ROIs | $F(2.145, 205.938) = 109.109$, $p < 0.001$, partial $\eta^2 = 0.532$ |
| | load x ROIs | $F(2.185, 209.765) = 1.518$, $p = 0.220$, partial $\eta^2 = 0.016$ |
| | AQ group x ROIs | $F(2.145, 205.938) = 0.295$, $p = 0.760$, partial $\eta^2 = 0.003$ |
| | load x AQ group x ROIs | $F(2.185, 209.765) = 0.592$, $p = 0.569$, partial $\eta^2 = 0.006$ |
| **70 Hz** | | |
| social | load | $F(1, 96) = 0.721$, $p = 0.398$, partial $\eta^2 = 0.007$ |
| | AQ group | $F(1, 96) = 0.584$, $p = 0.447$, partial $\eta^2 = 0.006$ |
| | load x AQ group | $F(1, 96) = 0.001$, $p = 0.970$, partial $\eta^2 = 0.000$ |
| | ROIs | $F(2.146, 206.060) = 108.085$, $p < 0.001$, partial $\eta^2 = 0.530$ |
| | load x ROIs | $F(2.153, 206.719) = 1.562$, $p = 0.211$, partial $\eta^2 = 0.016$ |
| | AQ group x ROIs | $F(2.146, 206.060) = 0.268$, $p = 0.780$, partial $\eta^2 = 0.003$ |
| | load x AQ group x ROIs | $F(2.153, 206.719) = 0.660$, $p = 0.529$, partial $\eta^2 = 0.007$ |

| | | |
|---|---|---|
| **70 Hz** | | |
| visual | load | $F(1, 96) = 2.055$, $p = 0.155$, partial $\eta^2 = 0.021$ |
| | AQ group | $F(1, 96) = 2.373$, $p = 0.127$, partial $\eta^2 = 0.024$ |
| | load x AQ group | $F(1, 96) = 0.341$, $p = 0.561$, partial $\eta^2 = 0.004$ |
| | ROIs | $F(1.925, 184.757) = 121.715$, $p < 0.001$, partial $\eta^2 = 0.559$ |
| | load x ROIs | $F(2.392, 229.638) = 0.719$, $p = 0.512$, partial $\eta^2 = 0.007$ |
| | AQ group x ROIs | $F(1.925, 184.757) = 0.875$, $p = 0.415$, partial $\eta^2 = 0.009$ |
| | load x AQ group x ROIs | $F(2.392, 229.638) = 0.673$, $p = 0.537$, partial $\eta^2 = 0.007$ |

**Table S9. Left Dorsolateral Prefrontal Cortex (lDLPFC).** The x, y, z coordinates are indicated for the left dorsolateral prefrontal cortex according to the Montreal Neurological Institute (MNI). We based the coordinates on Meyer et al. (2015, 2012).

| ROI name | ROI abbreviation | X (MNI) | Y (MNI) | Z (MNI) | Reference |
|---|---|---|---|---|---|
| left dorsolateral | lDLPFC | -45 | 17 | 28 | Meyer et al. (2012) |
| prefrontal | | -45 | 30 | 24 | Meyer et al. (2015) |
| cortex | | -51 | 9 | 39 | Meyer et al. (2015) |

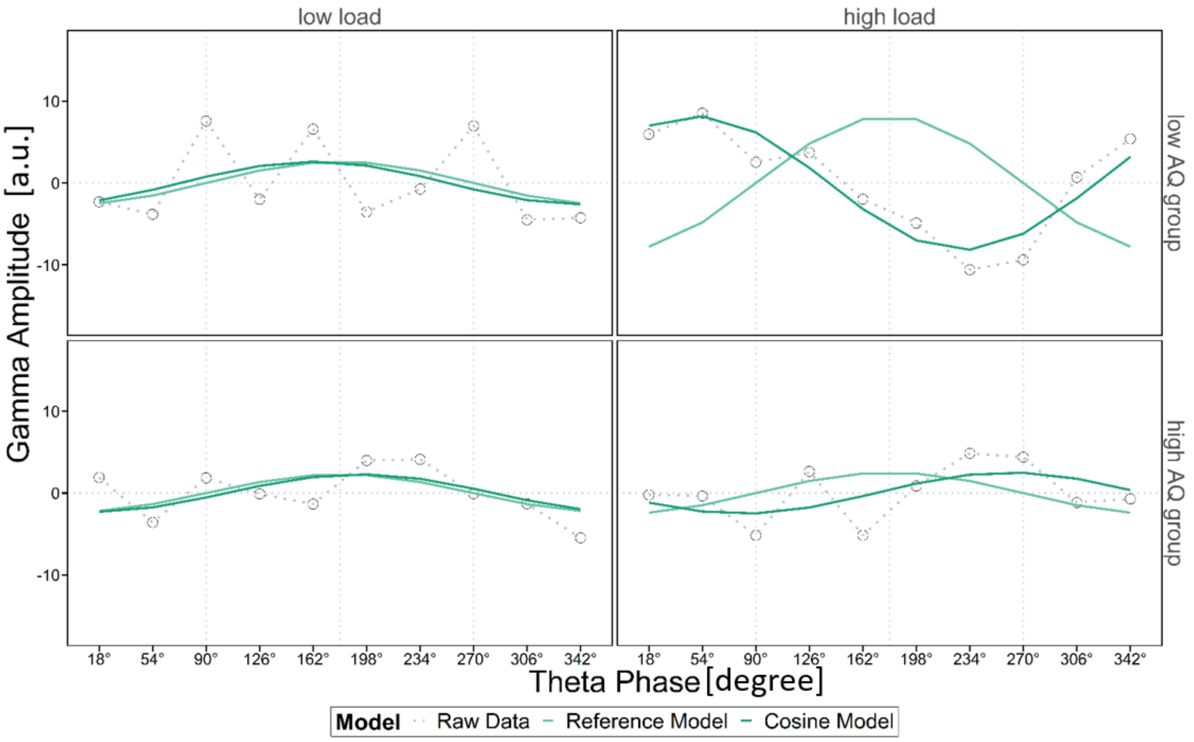


**Figure S7. Left DLPFC Phase-Amplitude Coupling in the Verbal Task.** Theta phase was extracted from the left dorsolateral prefrontal cortex (lDLPFC). 50-Hz posterior gamma amplitude was extracted from 11 posterior regions of interest (ROIs, see Table S7). The z-transformed posterior instantaneous gamma amplitude was sorted according to instantaneous lDLPFC-theta phase and averaged over all 11 posterior ROIs. In the line charts, the grey dots indicate the empirical z-transformed and sorted 50-Hz gamma amplitudes, the light turquoise lines our null-shift reference cosine model (simulating that strongest gamma amplitudes were locked in the trough of theta phase) and the dark turquoise lines the cosine model fitted to our empirical data.

In the verbal task in the 50-Hz frequency band in the low autistic-traits (AQ) group (top row) in the low load condition, the intercept model described the data better than a cosine model (AICc = -8.65). In the high load condition, the cosine model described the data significantly better than the intercept model (AICc = 9.90) but there was a significant phase shift, indicating that gamma amplitude was rather locked closer to the peak than the theta trough. In the high autistic-traits group (bottom row), the intercept model described the data better than the cosine model in both load conditions ($AICc_{low\ load}$ = -6.65; $AICc_{high\ load}$ = -6.74).

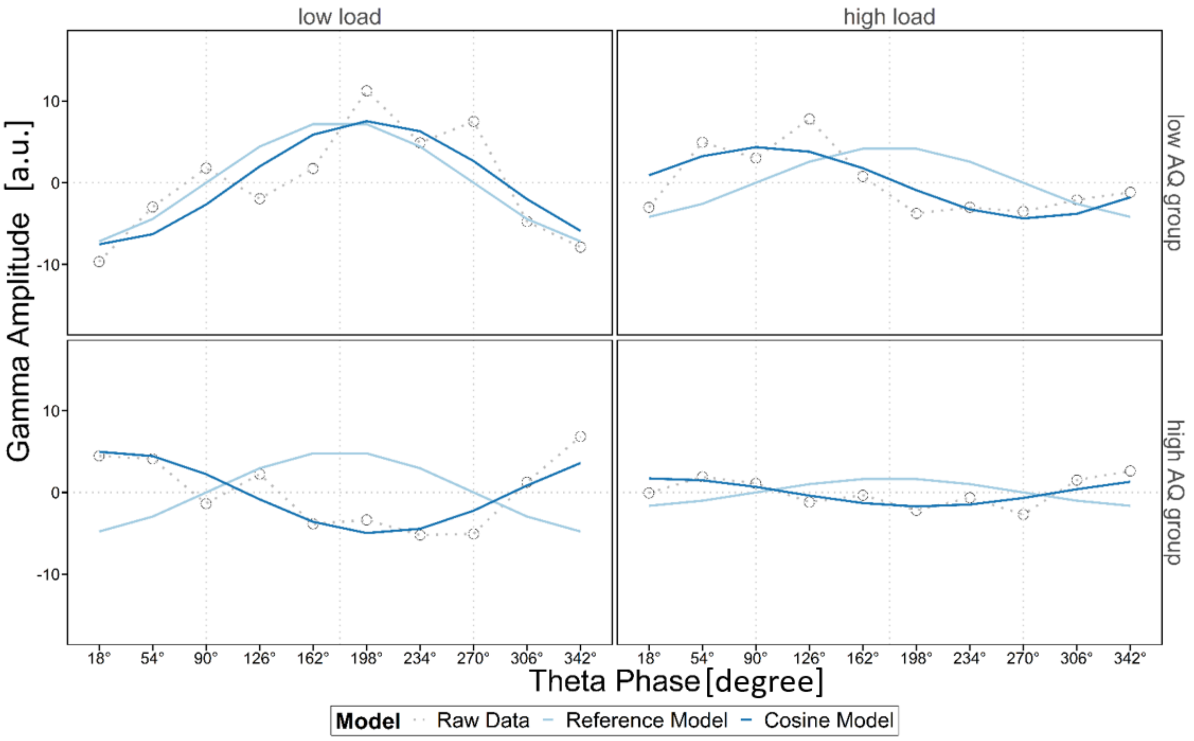


**Figure S8. Left DLPFC Phase-Amplitude Coupling in the Visual Task.** Theta phase was extracted from the left dorsolateral prefrontal cortex (lDLPFC). 50-Hz posterior gamma amplitude was extracted from 11 posterior regions of interest (ROIs, see Table S7). The z-transformed posterior instantaneous gamma amplitude was sorted according to instantaneous lDLPFC-theta phase and averaged over all 11 posterior ROIs. In the line charts, the grey dots indicate the empirical z-transformed and sorted 50-Hz gamma amplitudes, the light blue lines our null-shift reference cosine model (simulating that strongest gamma amplitudes were locked in the trough of theta phase) and the dark blue lines the cosine model fitted to our empirical data.

In the visual task in the 50-Hz frequency band in the low autistic-traits (AQ) group (top row), the cosine model did not significantly describe the data better than the intercept model ($AICc_{low\ load}$ = 1.92; $AICc_{high\ load}$ = 0.56). In the high autistic-traits group (bottom row) in the low load condition, the cosine model described the data significantly better than the intercept model (AICc = 3.10) and there was a significant phase shift, indicating that gamma amplitude was rather locked closer to the peak than the theta trough. In the high autistic-traits group in the high load condition, the intercept model described the data better than the cosine model (AICc = -2.71).
